# The modulatory role of early-life experiences and sex on cognitive resilience in APP/PS1 mice over time

**DOI:** 10.64898/2026.09.29.754953

**Authors:** Maria Carrigan, Jeniffer Sanguino-Gómez, Melis Bayer-Var, Olga Minko, Alise Bobkova, Eva Ruzicka, Laura López-Fedjajeva, Niek Renckens, Laurent Givalois, Paul J. Lucassen, Rik Ossenkoppele, Harm J. Krugers

## Abstract

**BACKGROUND:** Cognitive resilience to Alzheimer’s disease (AD) can be modelled in amyloid-expressing rodents through early handling (EH). This study investigated how EH and sex affect cognition and pathology over time.

**METHODS:** We assessed cognition (open field, object recognition test, Barnes maze) in 6- and 12-month-old APP/PS1 and WT mice (N=180, EH/Ctrl×M/F), performed biochemical and histological analyses of several brain areas to assess effects on Aβ, glial cells, and other markers of pathology, and investigated how behavior correlates with pathology.

**RESULTS:** EH improved non-spatial memory at 6 months, consistent with a reduction of hippocampal microglia density, and normalized locomotor activity at 12 months (both p<0.05). Females demonstrated similar (6 & 12 months) or better (12 months) cognition, despite greater pathology across timepoints. Correlations implicated microglial recruitment in EH-related effects, supported by morphological analysis of prefrontal regions (p<0.05).

**CONCLUSION:** Our findings highlight the critical role of early postnatal experiences and sex in shaping resilience in the context of AD pathology.

## BACKGROUND

Alzheimer’s disease (AD) is the leading cause of dementia, affecting millions of individuals worldwide each year, and posing an immense burden on patients, family members, caregivers and society as a whole [1]. Among the earliest pathological changes in AD is the excess accumulation of beta-amyloid (Aβ) which, in conjunction with a cascade of several molecular events, including neuroinflammatory changes, accelerates neurodegeneration and cognitive decline [2–5]. Interestingly, some individuals maintain their cognitive functions despite the presence of abundant Aβ proteinopathy; commonly referred to as cognitive resilience and defined as better than expected cognitive performance given a certain level of AD neuropathology [6,7].

Previous research has demonstrated that cognitive resilience can be modeled in transgenic Aβ-expressing rodents using the early-handling (EH) paradigm; a positive early-life intervention that can attenuate the progressive genotype-induced cognitive deficits otherwise observed in these AD mouse models [8–10], although the specific mechanisms underlying these behavioral effects remain unclear. Neuroinflammation has been proposed as another possible driver of neurodegeneration in AD, with the inflammatory response in the central nervous system primarily mediated by glial cells (astrocytes and microglia) exacerbating neuronal and synaptic loss [11]. While in the early stages of AD, glial cells may play a neuroprotective role [11,12], chronically active microglia around Aβ plaques transition to a pro-inflammatory phenotype, releasing inflammatory cytokines and losing their capacity for phagocytosis and lysosomal digestion, ultimately aggravating the disease [13–17]. Similarly, overactivation of astrocytes induces astrogliosis, which can also transform microglia to a reactive state, resulting in loss of their regulatory role, further contributing to inflammation and the dysregulation of the blood-brain barrier [18]. As such, variations in neuroinflammatory responses might underlie individual differences in cognitive trajectories in the face of AD neuropathology.

Interestingly, sex differences in AD have been established extensively, with females showing a higher baseline burden of neuropathology [19], faster cognitive decline [20,21] and more responsive immune systems compared to males [22–24]. Nonetheless, evidence suggests that females exhibit greater cognitive resilience [21,25] to AD, yet the biological basis of this advantage remains elusive. Thus, the present study aims to better understand the mechanisms underlying cognitive resilience, focusing on the modulatory role of positive early-life experiences and sex in a transgenic murine model of AD. Specifically, in a large-scale experimental setup, we exposed male and female mice expressing the APPswe/PS1dE9 (APP/PS1) genetic mutation and their wild-type (WT) littermates to either EH or control (Ctrl) conditions and assessed longitudinal cognitive performance across AD-relevant cognitive domains at 6 and 12 months of age, referencing earlier and later stages of human such as mild cognitive impairment and AD dementia, respectively. This resulted in a 2 × 2 × 2 design (Genotype: APP/PS1 vs. WT; Condition: EH vs. Ctrl; Sex: Male vs. Female) at two timepoints. In addition, brain tissue was collected at 12 months of age, as well as in a representative behavior-independent sample at 6 months, to compare neuropathological changes with cognitive function.

## METHODS

### Mice and breeding

Mice were housed under standard conditions with temperature maintained between 20–22°C and humidity at 40–60%. Standard food and water were provided ad libitum under a 12/12-hour light cycle, with lights on at 8 a.m.. Male and female WT and APP/PS1 animals of 6 months of age were used and tested over time (6 and 12 months of age). To obtain experimental mice, one 8-week-old C57Bl/6J virgin WT female (Envigo, The Netherlands) and one heterozygous male APP mouse were housed together for 1 week in a Type II long cage with standard corn-based bedding (Bio Services, Uden, Netherlands), and paper cage enrichment, including wood wool sizzle material, to allow mating. Individually housed pregnant females received a square piece of cotton 5 × 5 cm to prepare a nest (BMI Someren, The Netherlands) and were monitored daily between 9 a.m. and 11 a.m. for the birth of pups, and the day of birth was considered the previous day. The EH paradigm was initiated from post-natal day (P) 2 to 9 (see below and Fig. 1) as described previously [8]. At P28, mice were weaned and ear tissue was collected for identification and genotyping by standard PCR analysis. Littermates were housed with 2–5 same-sex littermates per cage, irrespective of genotype. Single females were housed with females from other litters, whereas single males were excluded from all subsequent procedures. Experimental males were housed in a separate room according to standard housing practices to minimize reproductive-related stimulation associated with the presence of females. All animals were left undisturbed (except for cage cleaning once a week) until the start of the experimental procedures, as well as between timepoints. Group sizes were chosen to ensure sufficient statistical power. Final number of mice—males: WT-Ctrl, 16; APP/PS1-Ctrl, 15; WT-EH, 17; and APP/PS1-EH, 12; females: WT-Ctrl, 14; APP/PS1-Ctrl, 15; WT-EH, 17; and APP/PS1-EH, 14. In addition, littermates not selected for behavioral testing were sacrificed as described below at 6 months of age for post-mortem processing, representative of brain pathology levels at 6 months in the longitudinal cohort. These numbers were as follows; males: WT-Ctrl, 8; APP/PS1-Ctrl, 8; WT-EH, 7; and APP/PS1-EH, 8; females: WT-EH, 8; APP/PS1-Ctrl, 5; WT-EH, 8; and APP/PS1-EH, 8. All experimental procedures were performed in accordance with Dutch national law as well as under directive 2010/63/EU of the European Parliament on animal experiments. The animal welfare committee of the University of Amsterdam and the Central Authority for Scientific Procedures on Animals (CCD) approved all experiments.

**Figure 1.**
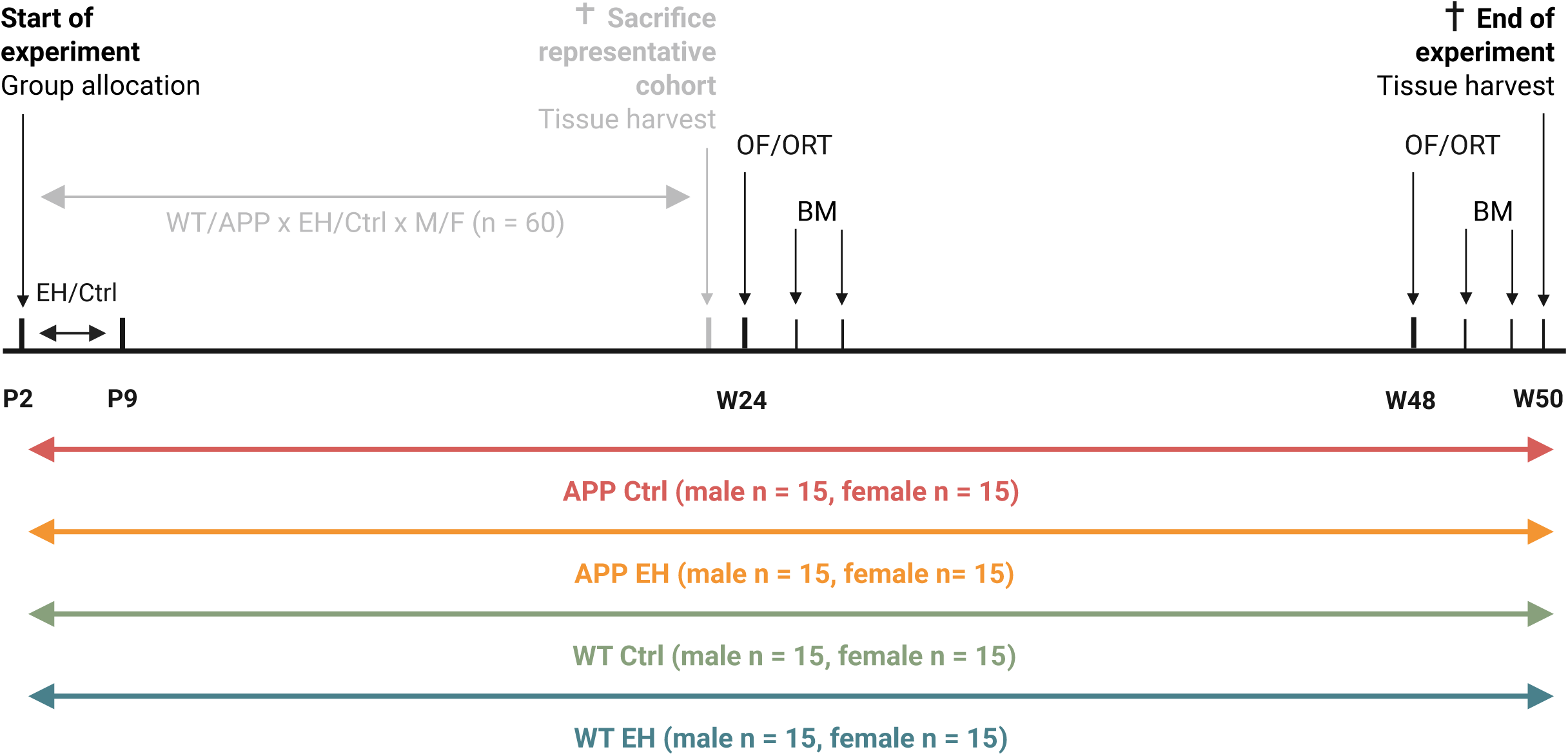
Study design. Experimental timeline of male and female APP/PS1 (APP) mice and wild-type (WT) littermates. At post-natal day (P) 2, assignment to early-handling (EH) or control (Ctrl) conditions, followed by brief daily maternal separation in the EH group from P2-9. Behavioral testing of the main cohort (colored groups, N = 120) at 6 (Week [W] 24) and 12 months (W48), including locomotor and anxiety-related behavior assessment in the open field (OF), non-spatial discrimination and memory in the novel object recognition test (ORT), and the Barnes maze (BM), assessing spatial learning, memory and cognitive flexibility. Tissue was harvested after behavioral acquisition at 12 months. In a separate cohort (grey group, N = 60) tissue was collected at 6 months. Numbers indicated for the main cohort represent intended group sizes; final group sizes ranged from 12–17 animals.

### Early handling

At P2, all litters were weighed and culled to six pups per litter, and dams and their litters were randomly assigned to the EH or control (Ctrl) condition until P9, after which all mice were treated equally (Fig. 1). A balanced sex ratio was maintained whenever possible, with at least one pup of each sex per litter. Ctrl mice were housed with normal nesting and bedding material and were left undisturbed between P2-9. EH mice underwent a daily 15-minute maternal separation from P2-9 between 9 a.m. and 11 a.m. During separation, dams and pups were placed in clean separate cages and reunited after 15 minutes in their home cage. Each litter and dam were assigned to their respective separation cages on P2, and these cages remained constant throughout all EH procedures. Pup cages were kept on a 26 °C heating pad and lined with home-cage bedding to minimize stress by maintaining warmth and home-cage smell. On P9, all mice were weighed again and placed in standard cages, with sufficient bedding and nesting material until weaning at P28. In total, 23 litters were assigned to the Ctrl condition and 24 litters to the EH condition, resulting in 120 Ctrl and 143 EH mice. Animals for the main and representative cohorts were subsequently selected based on sex and genotype.

### Maternal behavior

Maternal behavior was observed daily for 15 minutes from P2-9 immediately upon reunion (i.e., following separation) in EH nests, and between 9 a.m. and 11 a.m. in Ctrl nests, to assess whether the EH paradigm increased maternal care. The level of activity of the dam was scored (frequency) in a number of behaviors, which can be classified in the following groups: licking and grooming behavior, nursing behavior, and the net contact time between the dam and the pups.

### Cognitive assessment

All behavioral tests were carried out between 9 a.m. and 5 p.m., and all animals were allowed to habituate to the testing environment for 30 minutes before each behavioral test. Males were always tested prior to the females to minimize potential olfactory bias associated with reproductive-related stimuli. All testing and performance assessments were carried out by an experimenter blinded to the experimental group. Testing order of animals, equipment, and trial setup were randomized where appropriate within each sex. Arenas and test equipment were cleaned with 20 % ethanol between each trial to eliminate odor cues. During all tests, mice were recorded using a video camera (Basler Cam 4, Baseler AG, Germany) connected to a computer utilizing Ethovision XT 16 software (Noldus, The Netherlands).

### Open field

APP and WT male and female mice were assessed for general locomotion and anxiety-like behavior using the open field test (OF) [26]. Mice were placed individually in a rectangular arena (50 × 27.5 cm) with high walls and allowed to explore freely for 10 minutes. Locomotor activity was quantified by total distance travelled and movement velocity, while anxiety-related behavior was assessed via time (%) spent in the periphery compared to the center area.

### Novel Object Recognition

Non-spatial memory was assessed in the novel object recognition test (ORT) [27] over two consecutive days in a 50 × 27.5 cm open field. Mice were exposed to two identical objects for 5 minutes to familiarize them (training phase). After 24 h, one familiar object (FO) was replaced with a novel object (NO), and mice were allowed exploration for 5 minutes. Object exploration time and the Discrimination Index (DI = [Time around NO – Time around FO] / [Time around NO + Time around FO]) were calculated, with positive DI indicating preference for the NO.

### Barnes maze

Spatial learning, memory, and cognitive flexibility were tested in the Barnes maze [28]. Mice were tested in a standard circular maze (110 cm diameter) with 12 equidistant exit holes (5 cm diameter). The platform was elevated 153 cm above ground. Behavioral testing was initiated with a free exploration trial (FET), during which all exit holes remained closed, 24 h prior to the acquisition phase. During the acquisition phase (assessing spatial learning), animals were placed in the center of the maze twice daily with a 30-minute interval between trials, over four consecutive days, and allowed to find the exit hole leading to an escape box lined with a tissue and positioned below the maze. Upon escape, the mouse was left in the box undisturbed for 15 seconds before being returned to its home cage. If the mouse exited the box during this period, it was promptly removed from the platform and returned to its home cage. In case of failure to locate or enter the escape hole within three minutes, the mouse was gently guided to the correct location and was allowed to remain in the escape box for 30 seconds without interference. A probe trial assessing spatial memory, in which all exit holes were closed, was conducted 24 hours after the last acquisition trial. To assess behavioral flexibility, the exit hole was then moved to a new location (180° rotated) for a reversal learning phase lasting four days, followed by another probe trial. At 12 months, the FET was treated as an additional probe trial to assess long-term memory (SA5, supplementary material). To enhance motivation to escape, aversive stimuli were employed, including white noise (±30 dB) and a continuously running ventilator situated above the maze. In addition, light conditions were controlled, with an intensity of 0.65 lux when the light was off, and 40 lux when illuminated at the start of each trial. Distal visual cues outside the maze remained fixed relative to the exit hole.

### Tissue preparation

Following behavioral testing at 12 months (or at 6 months of age in the behavior-independent, age-representative cohort), mice were sacrificed by acute decapitation between 1 p.m. and 3 p.m. Brains were collected, of which the right hemispheres were preserved by immersion fixation in 4% paraformaldehyde solution in phosphate buffer (PB, 0.1 M PB, pH 7.4) for 24 h, followed by storage in a PB sodium azide (0.01%) solution at 4°C until further processing. The tissue was cryoprotected using sucrose solutions (15% sucrose/ 0.1 M PB solution for 24 h [or until sunken], then transferred to 30% sucrose/ 0.1 M PB solution 24 h). After cryoprotection, the hemispheres were snap frozen on dry ice and stored at −80°C until slicing; 40 µm thick sections were cut using a sliding microtome and stored in a PB-0.01% sodium azide solution at 4°C or anti-freeze solution (50% [v/v] 0.05 M PB-saline [PBS], 30% [v/v] ethylene glycol, and 20% [v/v] glycerol) at −20°C until immunohistochemical staining. From the left hemispheres, the hippocampus was isolated, flash-frozen in dry ice, and stored at −80°C for further processing.

### Immunofluorescent immunohistochemistry

A triple immunofluorescence staining was performed to detect Aβ plaques, microglia (IBA1) and reactive astrocytes (GFAP) in 40-µm-thick free-floating sections of the right hemisphere. Brain sections were washed 3 x 5 min in PBS containing 0.3% Triton X-100 (PBS-T), followed by 30 min incubation with Fab fragment goat anti-mouse IgG (H+L) (1:200, 115-007-003, RRID: AB_2338476, Jackson ImmunoResearch) to block endogenous mouse IgG. After 3 x 5-min washes in PBS-T, non-specific binding sites were blocked for 2 h at room temperature (RT) in a blocking buffer: PBS-T with 5% normal goat serum (NGS) and 5% normal donkey serum (NDS). Sections were then incubated overnight at 4°C with the primary antibody mix: PBS-T with 2% NGS and 2% NDS, rabbit anti-GFAP (1:10000; ab7260, Abcam), guinea pig anti-Iba1 (1:1000; 234 308, Synaptic Systems), and mouse anti-human Aβ (1:1500; SIG-3932-1000, BioLegend). The next day, sections were washed 3 x 5 min in PBS-T and incubated for 2h at RT while protected from ambient light with the secondary antibody mix: PBS-T with 1% NGS and 1% NDS, DAPI (1:1000; 10184322, Thermo Fisher), goat anti-rabbit Alexa 568 (1:500; A11011, Thermo Fisher), goat anti-guinea pig Alexa 647 (1:500; A21450, Thermo Fisher), and donkey anti-mouse Alexa 488 (1:500; A21202, Thermo Fisher). Finally, sections were washed 3 x 5 min in PBS and mounted onto slides in 0.1 M phosphate buffer (PB) and subsequently cover slipped with Fluoromount-G™ Mounting Medium (00-4958-02, Invitrogen).

### Imaging and quantification

The SLIDEVIEW VS200 universal whole slide imaging scanner (Evident Scientific, The Netherlands) was used to visualize the immunofluorescent signal in the 405nm (DAPI), 488nm (Aβ), 568nm (GFAP), and 647nm (IBA1) channels. Plaque load, the number of IBA1-and GFAP-positive cells, area coverage, as well as microglia morphology and plaque proximity of microglia, were quantified for all mice by an experimenter who was blind to the experimental conditions using Fiji (ImageJ). OME-TIFF hyper stacks were imported and converted to calibrated images with a pixel size of 0.3450 x 0.3450 µm. Regions of interest (ROIs) were selected based on their biological relevance in AD-dependent compromised cognitive function and defined using the DAPI channel (Ch1) to ensure the proper anatomical localization while remaining blind to the other marker expressions. For each selected brain region (dentate gyrus [DG], cornu ammonis 1 [CA1], prelimbic area [PLA], anterior cingulate cortex [ACC], infralimbic area [ILA]), a single square ROI of 250 x 250 µm was placed manually by positioning the central landmark point on the DAPI channel. All ROIs were selected based on the Allen Coronal Mouse Atlas. For the prefrontal subregions (PLA, ACC, ILA; roughly corresponding to Bregma range of 2.96 mm to 1.70 mm), the landmark was always placed onto the layers II/III of each region, whereas for the hippocampal (approximately Bregma range −1.22 mm to −3.80 mm) subregions (DG, CA1), the molecular layer of the DG and the stratum radiatum layer of the CA1 were used. Based on the image calibration, the size of the square ROI corresponded to 725 x 725 pixels. The number of detected plaques or cells within the ROI was divided by the ROI size (62,500 µm²) and then multiplied by 1,000,000 to convert counts to density per mm² for statistical analysis. Marker area coverage was calculated as the percentage of the total ROI area occupied by the positive signal using intensity thresholding. For microglia morphology and plaque proximity of microglial cells, acquired ROI images were selected based on the Allen Mouse atlas, roughly within Bregma of 2.22 mm to 1.70 mm for prefrontal subregions and Bregma range −1.22 mm to −1.94 mm for hippocampal subregions, and subdivided into 5 images of 500 × 500 pixel per ROI (four corners plus a center crop), from which the most central cell was selected for further processing with Microscopic Image Analyzer Threshold (MIAThreshold) software. Matched Aβ/Iba1 TIFF images were analyzed in Fiji using a custom macro. The Aβ channel was smoothed using a Gaussian blur (σ = 1), background was subtracted using a rolling-ball radius of 50 pixels, and plaques were segmented using the Auto Local Threshold (Huang) method. Candidate plaques were identified by particle analysis using a minimum size of 50 pixels and circularity of 0.20–1.00; elongated objects (aspect ratio > 1.5) were excluded. For each detected plaque, a donut-shaped ROI was generated to define the plaque vicinity, using a peripheral band with a width equal to one-twelfth of the mean plaque width and height, while excluding the plaque core. Iba1-positive microglia located within this plaque-associated ROI were classified as plaque-associated microglia (PAM), whereas cells outside the ROI were classified as non-plaque-associated microglia (NPAM).

### SDS-soluble and −insoluble Aβ levels

Aβ42 peptide levels were determined in whole hippocampal homogenates extracted from the left hemispheres using a sandwich enzyme-linked immunosorbent assay (ELISA; Human Aβ42 Ultrasensitive ELISA Kit, Invitrogen). In brief, brain tissue was homogenized with extraction buffer by sonication (5 & 10 seconds on ice) before ultracentrifugation at 14000 rpm for 30 minutes at 4 °C. Obtained supernatants were used to quantify soluble fractions of Aβ42 peptides. Insoluble fractions were further processed by homogenization with sonication (5 & 10 seconds on ice) under addition of 70% formic acid before ultracentrifugation at 14 000 rpm for 30 min at 4°C. Obtained supernatant was supplemented with Trisma base 1.5 M to quantify insoluble fractions. Results were normalized to protein concentrations calculated using the bicinchoninic acid method (BCA kit, Pierce) for soluble fractions, and the NanoDrop Spectrophotometer (Thermofisher Scientific) for insoluble fractions.

### Statistical analysis

Statistical analysis was performed in R version 4.3.2 (The R Foundation, Vienna, Austria). Barnes maze data were preprocessed prior to statistical analysis using in-house pipelines. Data were considered statistically significant when p < 0.05 and are expressed as mean ± standard error of the mean in figures and mean ± standard deviation in the text. Animals/observations were excluded in case of technical failures, following spontaneous death (see also **supplementary Material, SF1 & SA1**) and/or if identified as an extreme outlier using the IQR-method (3 × IQR). Between-group comparisons and interaction terms were assessed using linear (mixed) models with F-tests stratified by genotype, unless otherwise specified. Behavioral analysis was corrected for total distance traveled where appropriate to account for differences in activity levels between groups. Animal identity was added as random factor in repeated measures analyses (Barnes maze learning curve, longitudinal analyses). One-sided Student’s t-tests were performed in select groups to assess preference (NO vs. FO in the ORT) against zero and to compare performance on the probe trials of the Barnes maze against chance level (8.3%). Postmortem data were analyzed per subregion (DG, CA1, ACC, PLA, ILA). In addition, we calculated hippocampal and prefrontal composites (HIP and PFC, respectively) by averaging across the respective subregions. Sensitivity analyses were performed to account for effects of litter or staining batch where appropriate. Pearson’s correlation coefficients to study behavior-pathology associations were calculated separately for each sex and condition subgroup. Behavioral parameters were flipped where appropriate so that higher scores reflect better performance across all variables. Linear regression models including sex or condition interaction terms were used to test whether the associations between variables differed by sex or condition. Microglia morphology was analyzed using a Sholl analysis with quantification of area under the curve (AUC) to measure branching complexity. In addition, hierarchical clustering on principal components (HCPC) was performed to assess differences in microglia morphology subpopulations (morphotypes). Prior to cell-level clustering, which was performed across all data (i.e., both genotypes) with animal identity added as a random factor, each morphological metric was normalized and z-transformed to center the data. A principal component analysis (PCA) was conducted on PCs which had an eigenvalue > 1. The relative distribution of morphological clusters per conditions was evaluated in all regions and composites. Significant differences in cluster distribution per condition were estimated using a global Permutational Multivariate Analysis of Variance (PERMANOVA) with FDR (Benjamini-Hochberg) correction across pairwise comparisons. In addition, due to insufficient staining quality in a subset of 6-month animals (n = 9), these samples were re-stained. To correct for resulting batch-effects, batch 2 measurements were multiplicatively scaled to the corresponding batch 1 means within each biological group (region × condition × genotype × sex), with correction factors estimated from animal-level means and batch 1 used as the reference.

## 3. RESULTS

### 3.1 Maternal care

#### EH increased maternal care for the offspring

Mice were subjected to EH or Ctrl conditions during P2-9. EH dams showed increased maternal behavior across active measures including arched-back nursing, licking/grooming and nest building (**SA2)**, consistent with previous reports [8]. This did not result in body weight differences between EH or Ctrl litters (**SA3**).

### 3.2 Cognitive performance in 6- and 12-month-old APP/PS1 mice

#### 3.2.1 EH reduces APP/PS1-induced changes in general locomotion at 12 months of age

To assess whether EH affected general locomotion and anxiety-like behavior, mice were tested in the Open Field Test at 6 and 12 months of age (**Fig. 2A**). At 6 months, there were no differences in general locomotion as measured by total distance traveled (cm). At 12 months, EH reduced genotype-induced hyperactivity (Ctrl: 3812 ± 1554 cm; EH: 2870 ± 1301 cm; F(1,29) = 4.25, p < 0.05 [**Fig. 2D**]); SF4 and SA4, supplementary material), and male mice traveled less than female mice (Male: 2778 ± 752 cm; Female: 4389 ± 1859 cm; F(1,29) = 9.95, p < 0.01 [Fig. 2E]), with no interaction effect. There were no differences in anxiety-like behavior, assessed by proportion of time spent in the periphery (%), at either 6 months or 12 months. Longitudinally, overall locomotor activity declined (6m: 50288 ± 3413 cm; 12m: 3413 ± 1507 cm; F(1,29) = 31.46, p < 0.001), consistent with increased periphery time between 6 and 12 months (6m: 63 ± 15 %; 12m: 69 ± 20 %; F(1,29) = 7.33, p < 0.05).

**Figure 2.**
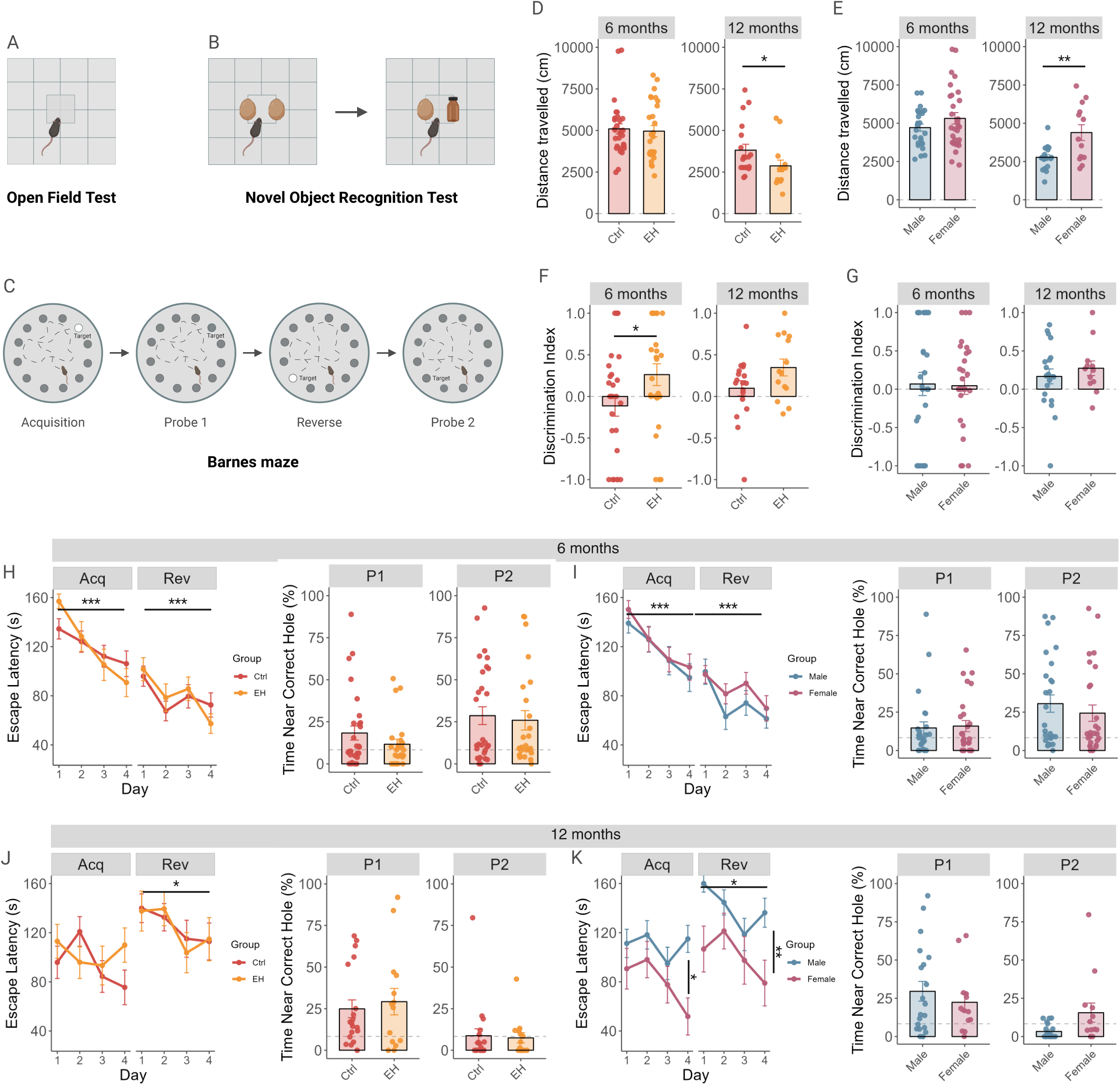
Behavioral outcomes by condition and sex in APP/PS1 mice. A) Open field test (OF). B) Novel object recognition test (ORT). C) Barnes maze (BM). D) EH reduces general locomotion (p<0.05) and E) females show hyperactivity compared to males (p<0.01) at 12 months in the OF. F) EH mice show better non-spatial memory at 6 months in the ORT (p<0.05), independent of sex (G). H-I) No group differences in spatial learning (Acquisition [Acq]; day effect: p<0.001), cognitive flexibility (Reverse acquisition [Rev]; day effect: p<0.001), or spatial memory (Probe [P] 1 & P2) in the BM at 6 months of age. J) EH did not alter Acq, cognitive flexibility (Rev, day effect: p<0.05) or memory at 12 months. K) Females show faster escapes (Acq, sex effect: p<0.05) and better cognitive flexibility (Rev, day effect: p<0.001; sex effect: p<0.01) trial, but no difference in spatial memory.

#### 3.2.2 EH affects non-spatial memory in 6- and 12-month-old mice

We assessed nonspatial recognition memory in the ORT at 6 and 12 months of age (Fig. 2B) as measured by the DI. At 6 months, Ctrl mice did not distinguish between the objects (−0.11 ± 0.68; t(29) = −0.93, p = 0.36), while EH animals preferred the novel object at borderline significance (0.26 ± 0.68; t(24) = 1.99, p = 0.06; condition effect: F(1,50) = 4.24, p < 0.05 [Fig. 2F]). Consistent with the beneficial effect observed at 6 months, at 12 months, animals that underwent EH showed a significant preference for the NO (0.35 ± 0.38; t(13) = 3.42, p < 0.01), as opposed to Ctrl mice, who failed to discriminate (0.10 ± 0.39; t(17) = 1.08, p = 0.29), albeit without a significant between-group improvement (condition effect: F(1,27) = 3.20, p = 0.08 [Fig. 2F]). We found no effect of sex (Fig. 2G), nor any interaction effects, on non-spatial memory among APP/PS1 mice at either age, and longitudinal analyses showed no significant group differences over time.

#### 3.2.3 At 12 months, females show better spatial navigation and cognitive flexibility than males

Spatial navigation, memory, and cognitive flexibility at 6 and 12 months of age were tested in the Barnes maze (Fig. 2C), measured by latency to escape (s) and time around the correct hole (%) during (reverse) acquisition and probe trials, respectively. Among APP/PS1 mice, all groups learned equally well during acquisition (day (D) effect: F(3,146) = 10.66, p < 0.001 [Fig. 2H-I]; D1: 145 ± 40 s; D2: 126 ± 53 s; D3: 109 ± 55 s; D4: 99 ± 57 s) and reverse training (day effect: F(3,146) = 6.39, p < 0.001 [Fig. 2H-I]; D1: 99 ± 45 s; D2: 73 ± 49 s; D3: 82 ± 49 s; D4: 66 ± 48 s) at 6 months of age, and there were no group differences in spatial memory in either probe trial. At 12 months of age, female mice outperformed males in both acquisition (sex effect: F(1,29) = 0.42, p < 0.05 [Fig. 2K]; Male: 110 ± 53 s; Female: 80 ± 57 s; day effect: F(3,86) = 2.23, p = 0.09) and reverse training (sex effect: F(1,29) = 7.76, p < 0.01 [Fig. 2K]; Male: 140 ± 50 s; Female: 101 ± 64 s; day effect: F(3,86) = 3.41, p < 0.05; D1: 139 ± 54 s; D2: 135 ± 49 s; D3: 110 ± 63 s; D4: 114 ± 64 s).

We found no other group differences in spatial learning or memory among APP/PS1 mice at 12 months (Fig. 2J).

### 3.3 Markers of pathology in 6- and 12-month-old APP/PS1 mice

#### 3.3.1 Females have higher levels of hippocampal Aβ in both age groups compared to males

To investigate whether EH and sex affected amyloid pathology (Fig. 3A, C-F) whether they interacted to influence amyloidosis, we first assessed whole hippocampal soluble and insoluble Aβ levels. We found that females had significantly higher levels of soluble Aβ levels at 6 (sex effect: F(1,25) = 12.50, p < 0.01; Male: 0.13 ± 0.06 mg/mL; Female: 0.25 ± 0.14 mg/mL) and 12 months (sex effect: F(1,29) = 9.58, p < 0.01 [Fig. 3A&D]; Male: 0.77 ± 0.35 mg/mL; Female: 1.17 ± 0.41 mg/mL), compared to males. Similarly, insoluble Aβ was higher in females at both 6 and 12 months, compared to males (6m: F(1,25) = 9.60, p < 0.01; Male: 44.00 ± 23.54 mg/mL; Female: 83.30 ± 47.10 mg/mL; 12m: (F(1,29) = 6.88, p < 0.05 [Fig. 3F]; Male: 277.36 ± 133.18 mg/mL; Female: 402.63 ± 134.75 mg/mL).

**Figure 3.**
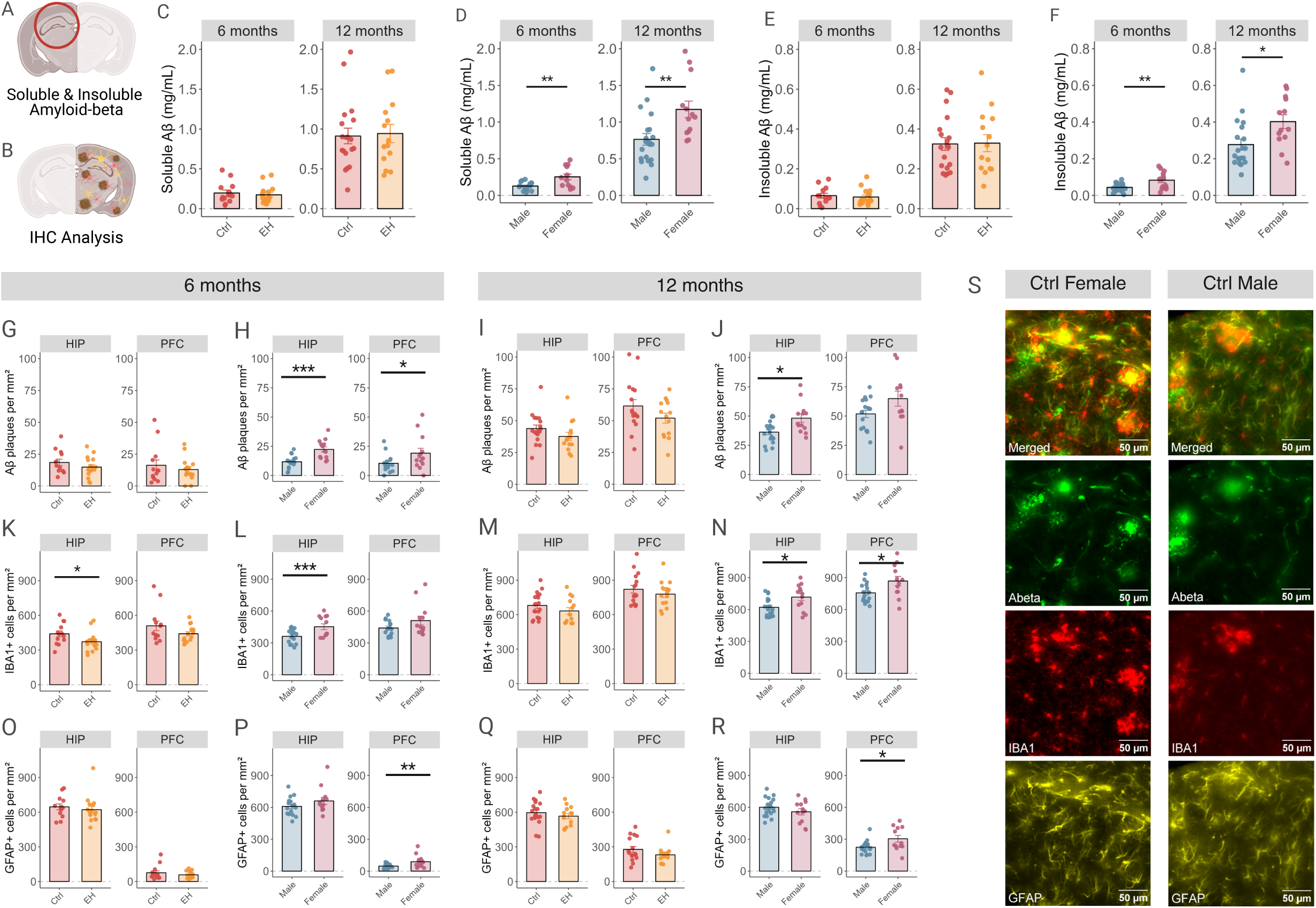
Markers of pathology in APP/PS1 mice by condition and sex. A) Whole-hippocampal measurements of soluble and insoluble Aβ. B) Immunohistochemical (IHC) staining for Aβ, microglia (IBA1), and reactive astrocytes (GFAP) was performed in the right hemisphere. C) EH did not alter soluble and (E) insoluble Aβ levels. D) Females show higher levels of soluble (6 & 12 months: p<0.01) and (F) insoluble Aβ (6 months: p<0.01, 12 months: p<0.05). G) EH did not affect Aβ-plaque density in the composites (HIP and PFC) at 6 and (I) 12 months. H) Females show higher hippocampal and prefrontal Aβ-plaque density at 6 months (HIP, p<0.001; PFC, p<0.05) and (J) in the hippocampus at 12 months (HIP, p<0.05; CA1, p<0.01; ILA, p<0.05). K) EH reduced IBA1+ cell density in the hippocampal composite (HIP, p<0.05) at 6 months, and (L) females have higher IBA1+ cell density (HIP, p<0.001). M) EH did not affect IBA1+ cell density in the composites at 12 months. N) Females show higher IBA1+ cell density in hippocampal (HIP, p<0.05) and prefrontal (PFC, p<0.05) regions at 12 months. O) No effect of EH on GFAP+ cell density in the composites at 6 or (Q) 12 months of age, whereas (P) females show higher prefrontal GFAP+ cell density (PFC, p<0.01) at 6 and (R) 12 months of age (PFC, p<0.05). S) Representative image of the Aβ-IBA1-GFAP triple staining, with the composite, Aβ, IBA1, and GFAP micrographs from top to bottom.

#### 3.3.2 EH affects Aβ plaques in the DG at 6 months, and females have higher plaque load

Complementing our assessment of soluble and insoluble Aβ levels, we investigated the potential effects of EH and sex, as well as their interaction, on Aβ-plaque density. Immunohistochemical analysis in the right hemisphere (Fig. 3B) revealed that at 6 months, Aβ-plaque density (plaques per mm^2^) in the DG was significantly lower among EH animals (condition effect: F(1,25) = 4.28, p < 0.05; EH: 22 ± 10 plaques/mm^2^; Ctrl: 28 ± 13 plaques/mm^2^ [SF2A]; HIP: p > 0.05 [Fig. 3G]), and females showed higher Aβ plaque burden compared to males in both hippocampal (HIP: F(1,25) = 21.31, p < 0.001; Male: 12 ± 6 plaques/mm^2^; Female: 22 ± 8 plaques/mm^2^; DG: F(1,25) = 19.95, p < 0.001; Male: 19 ± 9 plaques/mm^2^; Female: 32 ± 11 plaques/mm^2^; CA1: F(1,25) = 11.20, p < 0.01; Male: 5 ± 4 plaques/mm^2^; Female: 12 ± 8 plaques/mm^2^) and prefrontal regions (PFC: F(1,24) = 5.24, p < 0.05; Male: 10 ± 8 plaques/mm^2^; Female: 19 ± 15 plaques/mm^2^; ACC: F(1,23) = 5.97, p < 0.05; Male: 12 ± 13 plaques/mm^2^; Female: 26 ± 17 plaques/mm^2^ [Fig. 3H and Fig. 3S; SF3A]). Interestingly, female mice that underwent EH showed a significant reduction in Aβ-plaque density in the PFC compared to Ctrl mice (interaction effect: F(1,24) = 4.63, p < 0.05; post-hoc: p < 0.05; EH Female: 14 ± 10 plaques/mm^2^; Ctrl Female: 28 ± 18 plaques/mm^2^). At 12 months, Aβ-plaque density was not altered by EH in either ROI (Fig. 3I; SF2B). However, females showed higher plaque burden in the ILA (F(1,26) = 5.98, p < 0.05; Male: 40 ± 16 plaques/mm^2^; Female: 61 ± 26 plaques/mm^2^), as well as hippocampal areas (HIP: F(1,28) = 7.43, p < 0.05; Male: 36 ± 9 plaques/mm^2^; Female: 48 ± 13 plaques/mm^2^; CA1: F(1,28) = 8.49, p < 0.01; Male: 22 ± 12 plaques/mm^2^; Female: 38 ± 17 plaques/mm^2^ [Fig. 3J; SFB]), compared to males.

#### 3.3.3 EH reduced hippocampal IBA1+ cell density in 6-month-old mice

Given the prominent role of neuroinflammatory processes, we also investigated possible main and interaction effects of EH and sex on microglia. Assessment of IBA1 positive cells per mm^2^ revealed that among APP/PS1 mice, EH reduced IBA1+ cell density in hippocampal regions at 6 months (HIP: F(1,25) = 6.67, p < 0.05; EH: 372 ± 80 cells/mm^2^; Ctrl: 440 ± 92 cells/mm^2^ DG: F(1,25) = 8.73, p < 0.01; EH: 433 ± 103 cells/mm^2^; Ctrl: 531 ± 122 cells/mm^2^ [Fig. 3K; SF2C]), as well as non-significantly in the DG at 12 months (F(1,28) = 3.42, p = 0.07 [SF2D]). In addition, female APP animals showed higher levels of IBA1+ cells in hippocampal regions at 6 (HIP: F(1,25) = 13.95, p < 0.001; Male: 362 ± 64 cells/mm^2^; Female: 452 ± 96 cells/mm^2^; DG: F(1,25) = 17.87, p < 0.001; Male: 420 ± 88 cells/mm^2^; Female: 548 ± 121 cells/mm^2^; CA1: F(1,25) = 6.48, p < 0.05; Male: 304 ± 49 cells/mm^2^; Female: 357 ± 77 cells/mm^2^ [Fig. 3L; SF3C]) and 12 months (HIP: F(1,28) = 5.71, p < 0.05; Male: 620 ± 84 cells/mm^2^; Female: 716 ± 119 cells/mm^2^; CA1: F(1,28) = 7.41, p < 0.05; Male: 465 ± 90 cells/mm^2^; Female: 588 ± 151 cells/mm^2^ [Fig. 3N; SF3D]), as well as in prefrontal regions at 6 months (PFC: F(1,24) = 4.01, p = 0.06; Male: 440 ± 71 cells/mm^2^; Female: 510 ± 149 cells/mm^2^; ACC: F(1,23) = 9.10, p < 0.01; Male: 433 ± 94 cells/mm^2^; Female: 582 ± 180 cells/mm^2^ [Fig. 3L; SF3C]) and 12 months (PFC: F(1,26) = 5.47, p < 0.05; Male: 757 ± 84 cells/mm^2^; Female: 868 ± 152 cells/mm^2^; ILA: F(1,26) = 13.74, p < 0.01; Male: 644 ± 111 cells/mm^2^; Female: 810 ± 135 cells/mm^2^ [Fig. 3N; SF3D]). We found no other effects on IBA1+ cell density among APP/PS1 mice at either timepoint (Fig. 3M).

#### 3.3.4 Females show an increase in genotype-specific prefrontal GFAP+ cell densities

In addition to microglia, we investigated potential differences in reactive astrocytes in a similar statistical design, focusing on GFAP+ cell density in 6- and 12-month-old animals (Fig. 3O-R; SF2 and SF3). Females showed higher GFAP+ cell density in prefrontal regions at 6 months (PFC: F(1,24) = 8.62, p < 0.01; Male: 48 ± 21 cells/mm^2^; Female: 89 ± 59 cells/mm^2^; ACC: F(1,23) = 10.46, p < 0.01; Male: 45 ± 26 cells/mm^2^; Female: 112 ± 87 cells/mm^2^; ILA: F(1,22) = 3.70, p = 0.07; Male: 38 ± 26 cells/mm^2^; Female: 63 ± 37 cells/mm^2^ [Fig. 3P; SF3E]), as well as a non-significant increase in the DG (DG: F(1,25) = 3.72, p = 0.07). Interestingly, WT animals presented with negligible amounts of GFAP+ cells in prefrontal regions, indicating APP/PS1-specificity of reactive astrocytes in these ROIs (SF5, supplementary material). At 12 months, females had a higher GFAP+ cell density in prefrontal regions (PFC: F(1,26) = 5.03, p < 0.05; Male: 224 ± 60 cells/mm^2^; Female: 304 ± 110 cells/mm^2^; ILA: F(1,26) = 4.23, p < 0.05; Male: 206 ± 83 cells/mm^2^; Female: 298 ± 148 cells/mm^2^; PLA: F(1,26) = 3.38, p = 0.08 [Fig. Q, SF3F]), whereas males had a non-significant increase in the DG (F(1,28) = 3.20, p = 0.08). We found no other effects on GFAP+ cells among APP/PS1 animals.

### 3.4 Associations of cognition and pathology in 12-month-old APP/PS1 mice

#### 3.4.1 Spatial learning and memory associate with lower IBA1+ cell density among females

To assess whether and how differences in behavioral performance might be reflected by brain pathology markers, we ran exploratory correlation analyses at 12 months of age stratified by condition × sex. Among Ctrl females (Fig. 4A and SF6A), better cognitive flexibility as measured in the reverse acquisition phase of the BM associated with lower IBA1+ cell density (HIP: r = −0.8, p < 0.01; CA1: r = −0.67, p < 0.05; DG: r = −0.86, p < 0.001; PFC: r = −0.85, p < 0.01; ILA: r = −0.77, p < 0.05; PLA: r = −0.78, p < 0.05). Moreover, lower IBA1+ cell density in the hippocampus and prefrontal subregions associated with better spatial memory in Probe 1 (HIP: r = −0.86, p < 0.01; CA1: r = −0.74, p < 0.05; DG: r = −0.91, p < 0.01; ILA: r = −0.71, p < 0.05; PLA: r = −0.78, p < 0.05), as well as in prefrontal regions with better spatial memory in Probe 2 (PFC: r = −0.77, p < 0.05; ILA: r = −0.78, p < 0.01; r =-0.9, p < 0.05). Interestingly, anxiety-related behavior as measured by periphery time associated with lower IBA1+ cell density in the ILA (r = −0.81, p < 0.05). Among EH females (Fig.4C and SF6C), better spatial learning (r = −0.97, p < 0.05) and cognitive flexibility (r = −0.99, p < 0.001), as well as better spatial memory in Probe 2 (r = −0.96, p < 0.05), associated with lower IBA1+ cell density in the DG. Less hyperactivity associated with lower IBA1+ cell density in the CA1 (r = −0.98, p < 0.05), and less anxiety-related behavior associated with lower Aβ plaque density in the PLA (r = −0.99, p < 0.01). However, none of these associations survived multiple-comparison correction.

**Figure 4.**
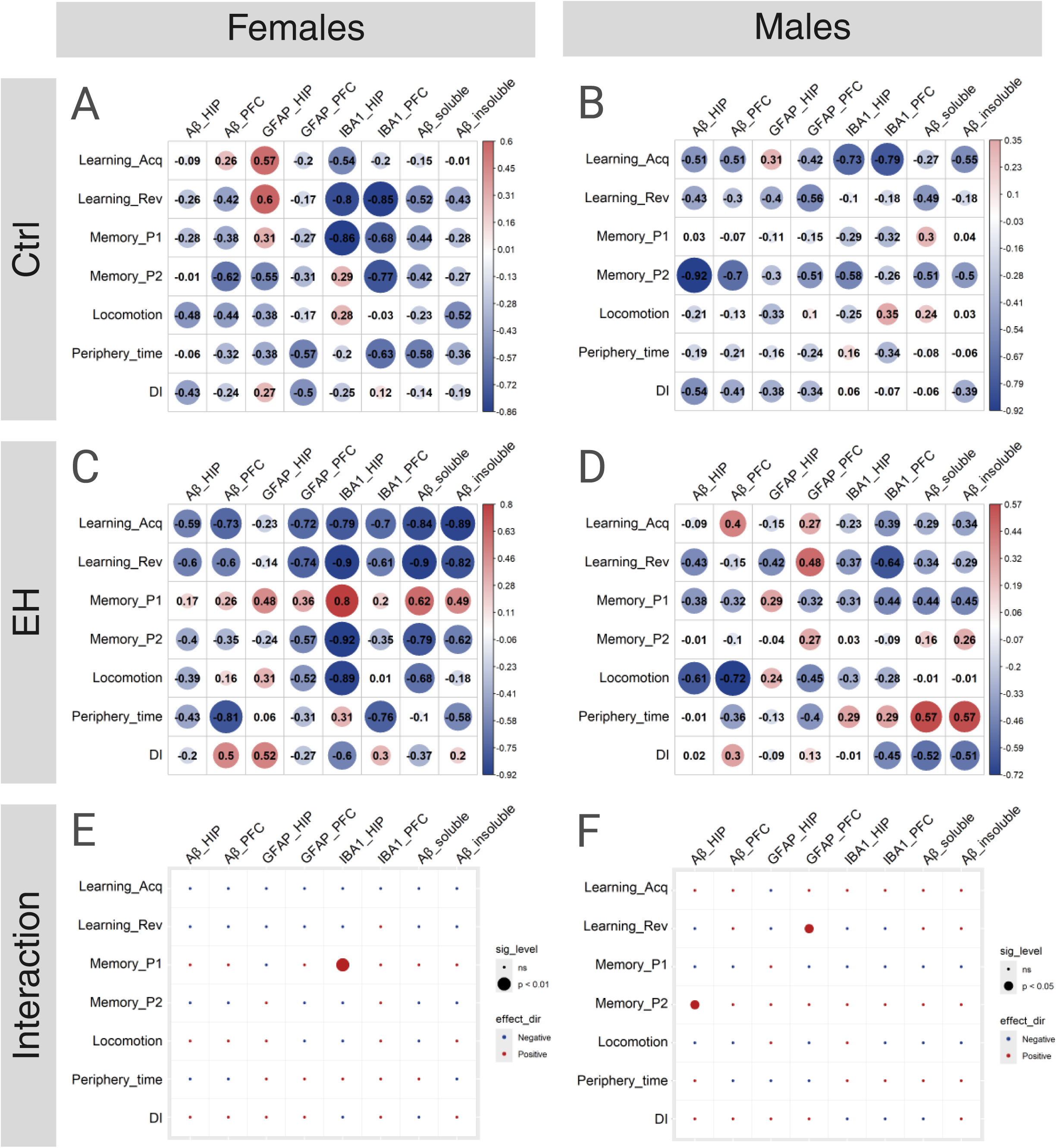
Associations of cognition and pathology in APP/PS1 mice at 12 months. Correlation analysis of behavioral parameters and measures of pathology (A-D) and sex-stratified interactions of condition and behavior-pathology associations (E-F). Higher scores on behavioral variables reflect better performance. A) Among Ctrl females, lower IBA1+ cell density was associated with better cognitive flexibility (reverse learning, HIP: r = −0.8, p < 0.01; PFC: r = −0.85, p < 0.01), better spatial memory in Probe 1 (HIP: r = −0.86, p < 0.01) and Probe 2 (PFC: r = −0.77, p < 0.05). B) Among Ctrl males, better spatial memory in Probe 2 associated with lower Aβ density in the HIP (r = −0.92, p < 0.001), and better spatial learning associated with lower IBA1+ cell density (HIP: r = −0.73, p < 0.05; PFC: r = −0.79, p < 0.05). C) Among EH females, there were no significant associations between behavior and markers of pathology in any of the composite regions. D) Among EH males, better cognitive flexibility associated with lower prefrontal IBA1+ cell density (PFC: r = −0.64, p < 0.05), and less hyperactive locomotion associated with lower prefrontal Aβ plaque density (PFC: r = −0.72, p < 0.05). E) We found a more positive association of better spatial memory in Probe 1 with higher hippocampal IBA1+ cell density in EH compared to Ctrl females (HIP: β = 0.23, SE = 0.06, t = 3.62, p < 0.01). F) Among males, EH had a more positive association between cognitive flexibility and prefrontal GFAP+ cell density (PFC β = 0.13, SE = 0.06, t = 2.19, p < 0.05), as well as between Probe 2 performance and higher hippocampal Aβ (HIP: β = 0.48, SE = 0.22, t = 2.17, p < 0.05).

#### 3.4.2 Spatial memory associates with lower Aβ plaque density among males

Among Ctrl males (Fig. 4B and SF6B), better spatial learning associated with lower Aβ density in the PLA (r = −0.81, p < 0.05), and better spatial memory in the Probe 2 associated with lower Aβ density in hippocampal regions (HIP: r = −0.92, p < 0.001, DG: r = −0.73, p < 0.05) and the PLA (r = −0.71, p < 0.05). In addition, spatial learning associated with lower IBA1+ cell density (HIP: r = −0.73, p < 0.05; DG: r =-0.78, p < 0.05; PFC: r = −0.79, p < 0.05; ILA: r =-0.71, p < 0.05), and Probe 2 performance associated with lower IBA1+ cell density in the DG (r = −0.69, p < 0.05). Interestingly, Probe 1 performance associated with higher GFAP+ cell density in the ILA (r = 0.89, p < 0.01). Furthermore, less anxiety-related behavior associated with lower IBA1+ cell density in the ILA (r = −0.74, p < 0.05). Among EH males (Fig. 4D and SF6D), better spatial learning associated with higher Aβ plaque density in the ILA (r = 0.7, p < 0.05), whereas better cognitive flexibility associated with lower Aβ plaque density in the CA1 (r = −0.71, p < 0.05) and lower IBA1+ cell density in the PFC (r = −0.64, p < 0.05). In addition, better Probe 1 performance associated with lower Aβ plaque density in the PLA (r = −0.74, p < 0.05), less hyperactive locomotion associated with lower Aβ plaque density in the PFC (r = −0.72, p < 0.05), and less anxiety-related behavior associated with lower GFAP+ cell density in the ILA (r = −0.63, p < 0.05). However, none of these associations survived multiple-comparison correction.

#### 3.4.3 Spatial memory is more strongly associated with higher hippocampal IBA1 density in EH females

Given our sex-specific results, we were interested whether condition might moderate the behavior-pathology associations within each sex. We found that condition moderated behavior-pathology associations among females (Fig. 4E and SF6E), such that better spatial memory in Probe 1 was more strongly associated with higher hippocampal IBA1+ cell density in EH compared to Ctrl mice (HIP: β = 0.23, SE = 0.06, t = 3.62, p < 0.01; CA1: β = 0.15, SE = 0.06, t = 2.31, p < 0.05; DG: β = 0.29, SE = 0.06, t = 4.70, p < 0.01). These effects did not survive correction for multiple testing, and we found no other significant effects of condition on the behavior and pathology associations with condition among females.

#### 3.4.4 EH males show stronger associations between behavior and pathology compared to Ctrl

Among males (Fig. 4F and SF6F), the association between spatial learning and higher IBA1+ cell density was stronger in EH animals compared to Ctrl in the ILA (β = 0.02, SE = 0.01, t = 2.82, p < 0.05). Similarly, EH had a more positive association between cognitive flexibility and prefrontal GFAP+ cell density (PFC: β = 0.13, SE = 0.06, t = 2.19, p < 0.05; PLA: β = 0.09, SE = 0.04, t = 2.24, p < 0.05), as well as between Probe 2 performance and higher hippocampal Aβ (HIP: β = 0.48, SE = 0.22, t = 2.17, p < 0.05; DG: β = 0.42, SE = 0.02, t = 2.27, p < 0.05). Conversely, the association between Probe 1 performance and lower Aβ in the PLA (β = −1.99, SE = 0.73, t = −2.74, p < 0.05), and the association between lower hyperactivity with lower IBA1+ cell density in the ILA (β = −7.25, SE = 2.53, t = −2.87, p < 0.05) was weaker in EH animals compared to Ctrl. Interestingly, the association between better non-spatial memory with higher Aβ plaque density in the DG (β = 0.03, SE = 0.02, t = 2.24, p < 0.05), and higher GFAP+ cell density in the ILA (β = 0.00, SE = 0.00, t = 2.72, p < 0.05) was stronger in EH compared to Ctrl. These effects did not survive correction for multiple testing, and we found no other significant effects of condition on the behavior and pathology associations with condition among males.

### 3.5 Microglia morphology in APP/PS1 mice

#### 3.5.1 Genotype-specific effects of EH and sex on branching complexity at 12 months of age

Based on the observed effects of EH and sex on microglial density, as well as our correlations pointing to potential differences in microglial recruitment, particularly in females, we next investigated microglia morphology. As expected, microglial branching complexity, as measured using Sholl analysis-derived area under the curve (AUC; Fig. 5A), was significantly reduced in PAM compared to NPAM in all regions (Fig. 5C,F; ST1, SF7E,F).

**Figure 5.**
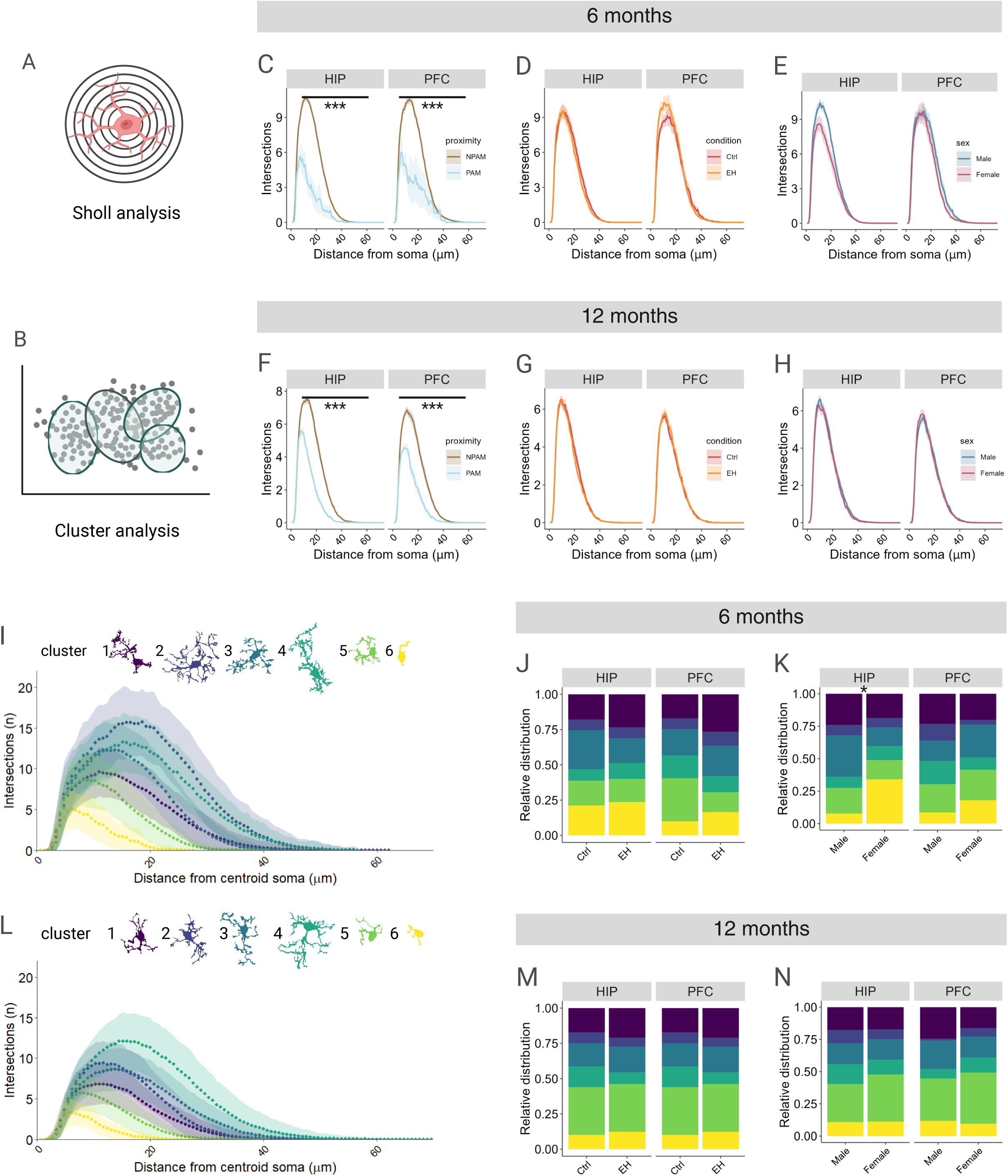
Microglia morphology at 6 and 12 months of age. Analysis of microglia morphology using A) Sholl analysis curves and B) Hierarchical clustering on principal components. C) Microglial branching complexity, as measured through area-under-the-curve (AUC) of Sholl curve, differed by plaque proximity at 6 months (HIP: F(1,506.78) = 28.05, p < 0.001; PFC: F(1,331.31) = 13.88, p < 0.001). D) No difference in AUC by condition or (E) sex at 6 months. F) Plaque proximity significantly affected AUC at 12 months (HIP: F(1,599.39) = 117.72, p < 0.001; PFC: F(1,424.44) = 71.63, p < 0.001), whereas no effects were found for (G) condition or (H) sex. I) Cluster morphology analysis at 6 months with representative cells for each cluster. J) Condition did not affect cluster distribution at 6 months of age. K) Females show a higher number of ameboid cells in hippocampal regions compared to males (F(1,32) = 6.99, p < 0.01). L) Cluster morphology analysis at 12 months with representative cells for each cluster. M) Condition and (N) sex did not affect cluster distribution at 12 months of age.

Interestingly, at 12 months of age, microglia of WT EH animals had a greater AUC in the hippocampus compared to WT Ctrl mice (HIP: F(1,56.46) = 6.03, p < 0.05; EH: 193 ± 93 intersections·µm; Ctrl: 164 ± 83 intersections·µm; DG: F(1,57.09) = 5.00, p < 0.05; EH: 191 ± 92 intersections·µm; Ctrl: 161 ± 83 intersections·µm; CA1: F(1,55.05) = 5.77, p < 0.05; EH: 195 ± 93 intersections·µm; Ctrl: 167 ± 83 intersections·µm [SF8B]), and microglia of WT females had significantly lower AUC than WT males in hippocampal regions (HIP: F(1,56.46) = 4.54, p < 0.05; Male: 193 ± 91 intersections·µm; Female: 165 ± 85 intersections·µm; CA1: F(1,55.05) = 6.03, p < 0.05; Male: 197 ± 91 intersections·µm; Female: 165 ± 85 intersections·µm) and the ILA (F(1,54.90) = 4.17, p < 0.05; Male: 200 ± 75 intersections·µm; Female: 174 ± 81 intersections·µm [SF8D]). We found no other effects of EH, sex, or interactions on AUC at either timepoint (Fig.5D,E,G,H; SF7A-D; SF8A,C).

#### 3.5.2 EH moderately affects cluster distribution in 12-month-old mice

As expected, cluster distribution (Fig.5B) differed significantly between PAM and NPAM in 6- and 12-month-old APP/PS1 mice (ST2). Moreover, at 6 months, hippocampal cluster distribution differed significantly by sex, with females showing less ramified cells compared to males (HIP: F(1,32) = 6.99, p < 0.01; cluster 3 post hoc: pFDR < 0.05; Male: 31.85 ± 22.65 %; Females: 14.25 ± 13.92 %; DG: F(1,31) = 3.80, p < 0.05; CA1: F(1,25) = 4.84, p < 0.001 [Fig. 5I,K; SF7I]). We found a similar trend for sex in prefrontal regions (PFC: F(1,23) = 2.32, p = 0.07; ACC: F(1,21) = 2.05, p = 0.08; ILA: F(1,19) = 2.31, p = 0.05), with a significant sex effect in the PLA (F(1,20) = 2.41, p < 0.05 [SF7I]). In addition, we found a significant effect of EH on cluster distribution in the ILA among APP/PS1 mice at 6 months (F(1,19) = 3.73, p < 0.01; cluster 5 post hoc: pFDR < 0.05; EH: 13.85 ± 28.73 %; Ctrl: 25.00 ± 21.64 %; cluster 6 post hoc: pFDR < 0.001; EH: 0.00 ± 0.00 %; Ctrl: 17.50 ± 28.24 % [SF7G]) and 12 months (F(1,36) = 3.23, p < 0.05 [SF7H]). Among WT animals, we found a significant effect of EH on cluster distribution in the PLA at 6 months (F(1,26) = 4.45, p < 0.01; cluster 1 post hoc: pFDR < 0.05; EH: 46.67 ± 19.52 %; Ctrl: 21.33 ± 24.46 % [SF8E]), and in hippocampal regions at 12 months (HIP: F(1,57) = 3.19, p < 0.05; cluster 4 post hoc: pFDR < 0.05; EH: 35.73 ± 22.42 %; Ctrl: 22.00 ± 14.13 %;DG: F(1,57) = 2.50, p = 0.06; CA1: F(1,57) = 2.28, p = 0.07 [SF8F]). We found no other effects of EH, sex, or any interactions on cluster distribution at either timepoint (Fig.5J,L-N; SF7H&J, SF8E-H).

## DISCUSSION

Despite abundant AD pathophysiology, some individuals maintain cognitive functioning and show only limited cognitive decline [6]. This study set out to elucidate the potential substrates underlying such cognitive resilience in the context of AD pathology by focusing on the role of early-life experiences and sex in a transgenic mouse model of AD. Specifically, we subjected male and female APP/PS1 mice and their WT littermates to EH, a positive early-life intervention previously shown to promote cognitive resilience [8], and characterized these mice cognitively and behaviorally at 6 and 12 months of age, reflecting early vs. late stages of AD progression in reference to human MCI and dementia stages, respectively. In addition, we assessed markers of AD pathology in brain tissue from these mice at 12 months of age, as well as in brain tissue from a representative 6-month-old cohort.

In line with previous research [8,10], we demonstrated that EH increased maternal care in their offspring, and found no negative effects of EH on pup body weight. At 6 months of age, we found no effects of EH or sex on genotype-induced hyperactivity in general locomotor behavior, whereas in EH APP/PS1 animals of both sexes at 12 months of age, general locomotion was normalized to WT levels. Our finding supports the previously described efficacy of early-life interventions to alter adult general locomotor behavior in the context of AD pathology [29], and demonstrate a beneficial effect of EH. Importantly, these effects were independent of anxiety-like behavior, as we found no differences in center versus periphery time across all animals at either timepoint. In contrast to a previous study [10], which found no effects of EH on object recognition memory at 4 months of age, we found better non-spatial memory in the ORT at 6 months of age among EH APP/PS1 mice relative to Ctrl APP/PS1 mice, suggesting that the beneficial effect of EH on non-spatial memory becomes evident in older APP/PS1 mice where dysfunction is more pronounced. In addition, although relatively high biological variability within groups may have occluded a treatment effect of EH at 12 months, EH APP/PS1 mice significantly preferred the novel over the familiar object, whereas Ctrl APP/PS1 mice failed to discriminate between the objects at this age. Sex did not moderate this effect, nor were there sex differences in object recognition memory at either age. Together, these findings suggest that EH indeed attenuates cognitive deficits in APP/PS1 mice irrespective of sex, however, in a seemingly progression-dependent manner given no previously described effects at 4 months [10]. Surprisingly, we found no effects of EH on spatial navigation, spatial memory, and cognitive flexibility at either timepoint, suggesting that the beneficial effect of EH – at least in the tasks used in our study - is limited to non-spatial cognitive function. Selective survival may also have contributed to the observed pattern, as our sensitivity analyses showed a higher hazard of death among EH APP/PS1 compared to Ctrl APP/PS1 mice, meaning that those surviving to 12 months may represent a subset with characteristics that differed from animals that did not survive, potentially reducing or modifying some of the effects of EH observed at earlier timepoints.

Interestingly, female mice outperformed males in both spatial navigation and cognitive flexibility at 12 months of age, supporting a female advantage in cognitive performance often seen in humans [25]. Such a female advantage in behavior is especially noteworthy in the context of the pathological states observed. That is, females displayed cognitive performance that was equal (ORT and BM at 6 months) or better (BM at 12 months) as compared to age-matched males, despite more advanced pathophysiology: We found higher levels of soluble and insoluble Aβ, higher overall Aβ plaque burden, higher microglia density, and higher prefrontal densities of reactive astrocytes among female APP/PS1 mice compared to males, supporting previous evidence of greater pathology and faster disease progression in females among mice [30] and humans [25]. Together, these findings suggest that female mice are more resilient to advanced pathophysiology compared to males, given similar or better cognitive performance despite higher pathological burden. Moreover, in support of such higher female resilience, prefrontal Aβ-plaque density was reduced in female EH APP/PS1 mice relative to female Ctrl APP/PS1 mice, whereas Aβ-plaque density in the DG was reduced by EH across sexes. In contrast, we found no effects of EH on microglia or astrocyte density in the CA1, despite the amyloid-beta accumulation and plaque load traditionally being prominent in this region, temporally followed by the DG [31]; although it is possible that we found no effects because pathology was already too advanced in this region, or because the effects of EH were too subtle relative to the extent of pathology. These findings also highlight the distinction between cognitive resilience and resistance to pathology. While preserved cognitive function despite pathology is consistent with resilience, the reductions in Aβ-plaque burden observed following EH may reflect an effect on the development or accumulation of pathology, and therefore a resistance-related process. However, we found no effects of EH on Aβ-plaque density in either hippocampal or cortical regions at 12 months of age, and the EH-induced hippocampal microglia density reduction at 6 months was no longer significant at 12 months of age. These findings suggest that with increasing age and at more advanced stages of disease, effects of EH might no longer be sufficiently prominent to alter pathology, although, as described earlier, selective survival may also have contributed. Of note, we found no evidence supporting a beneficial effect of EH on reactive astrocytes at either timepoint, suggesting that the mechanisms underlying resilience might be independent of astrocytic reactivity or astrogliosis.

To investigate how our behavioral observations are reflected by pathology directly, we ran a series of exploratory correlation analyses stratified by condition and sex. Because none of these findings survived correction for multiple comparisons, these results should be considered hypothesis-generating. As such, our main findings consistently pointed to an association between lower microglial density and better spatial cognition across conditions and sex. These results are consistent with other studies indicating a role of immune cells on spatial learning and memory [32,33]. Interestingly, we found that lower anxiety-related behavior associated with lower ILA-specific microglia density among Ctrl but not EH animals [32]. While the role of microglial activity in anxiety-like behaviors has been demonstrated extensively [34,35], it remains unclear how exactly the ILA in particular is involved in these predominantly amygdala-related processes, and why this association was absent in EH animals. A possible explanation could be that microglial density alone does not sufficiently reflect underlying functional processes, which might be different in EH and Ctrl animals. Supporting this notion, EH showed more positive associations between better spatial cognition and higher microglia density in hippocampal regions among females and in prefrontal regions among males. The differing associations between microglia density and cognition across EH and Ctrl conditions suggest that microglial density alone may be insufficient to capture the underlying microglial processes associated with cognitive function.

Based on these results, we investigated microglia morphology and confirmed an Aβ plaque-related reduction in microglial branching complexity, as well as significant differences in microglia morphotype distribution, in PAM compared to NPAM across age and sex in all regions [36,37]. In line with our microglia density results and correlation analyses, EH affected microglia morphotype distribution in various prefrontal subregions across age and genotype, and in hippocampal regions among 12-month-old WT mice. However, neither condition nor sex showed any relevant interactions with Aβ plaque proximity, suggesting that any potential effects of EH or sex are not Aβ plaque specific. These findings align with existing literature supporting the potential of enriching environmental interventions to modify microglial phenotypes even despite the presence of substantial amyloidosis [38–40]. In light of this, our results suggest that EH-related processes may delay age- and early pathology-related morphological changes, however, these effects are subtle and such processes might not be sufficient to overcome advanced disease-associated changes in microglia morphology. In addition, APP/PS1 females showed a greater proportion of amoeboid microglia in the hippocampus and prefrontal subregions at 6 months compared to males, and microglial branching complexity in those regions was significantly lower among 12-month-old WT females compared to males, supporting a sex-specific neuroimmune response across age and genotype [22,24].

Taken together, our findings suggest greater resilience in females, who showed similar or better cognitive performance, despite a higher pathological burden than males. Moreover, EH successfully modeled resilience by improving cognitive performance and delaying cognitive decline, particularly during early disease stages. Finally, our findings implicate neuroimmune pathways, particularly microglia, as potential mechanistic substrates underlying resilience.

This study has several limitations. First, due to practical limitations of our experimental design, this study did not include an independent replication cohort. Replication would have required an additional cohort of animals undergoing the same experimental procedures, which was not feasible within the scope and practical constraints of the present study. Nevertheless, the inclusion of a separate 6-month cohort allowed us to examine pathology at this stage of disease, providing complementary evidence alongside the behavioral findings at 6 months. However, the 6-month cohort used for postmortem analyses did not undergo behavioral testing and as such, we cannot exclude possible effects of training on better outcomes, nor could we test behavior-pathology associations at this timepoint. Similarly, such effects of training cannot be ruled out in the 12-month cohort, who have undergone behavioral testing twice, which might act as a form of environmental enrichment through cognitive stimulation and consequently affect brain pathology [39,40]. Moreover, given that the APP/PS1 model is quite vulnerable and a considerable number of animals drops out as aging progresses [41,42], our cohort of mice at 12 months of age therefore constitutes a “resilient” subgroup in terms of selective survival of the fittest, irrespective of experimental condition. As such, some effects of EH that emerged at 6 months of age might have been concealed at 12 months of age. Future research should therefore investigate earlier timepoints of intermediate-advanced pathology, for example at 8 or 10 months of age [43]. As noted previously, survival also affected our sample size and composition of the 12-month cohort, especially among EH APP/PS1 animals, despite its beneficial effects on cognition and pathology. Another practical consideration is that experimental males and females were housed in different rooms to minimize reproductive-related stimulation. Although this arrangement reflected standard housing practices, room-specific environmental factors, including exposure to sex-specific environmental cues, cannot be completely excluded as contributors to the observed sex differences. Finally, although statistical significance was assessed at the conventional threshold of p < 0.05, a more stringent threshold of p < 0.005 has been proposed [44]. Thus, findings with p values between 0.005 and 0.05 should be interpreted cautiously and in the context of the magnitude and consistency of the observed effects.

In addition, future research should investigate the immediate effects of EH at PND9 on both dams and pups. In particular, assessing glucocorticoid levels in dams and pups following maternal–pup separation could help determine whether EH induces acute physiological stress responses, including changes in corticosterone levels in maternal milk. Although speculative, such changes could represent a potential mechanism through which EH influences later cognitive outcomes of the offspring. Future studies should also examine synaptic structure and function, including synaptic proteins and AMPA receptor expression, as potential substrates underlying the observed effects of EH on memory. Similarly, future studies should look into epigenetic effects of EH to unravel potential (epi)genetic targets of resilience. In conclusion, our findings suggest greater resilience in females and demonstrate that EH can model resilience by improving cognitive performance and delaying cognitive decline, potentially through modulation of neuroimmune pathways, particularly microglial states.

## DECLARATIONS

### ETHICS APPROVAL AND CONSENT TO PARTICIPATE

All experimental procedures were performed in accordance with Dutch national law as well as under directive 2010/63/EU of the European Parliament on animal experiments. The animal welfare committee of the University of Amsterdam and the Central Authority for Scientific Procedures on Animals (CCD) approved all experiments.

### CONSENT FOR PUBLICATION

Not applicable

### AVAILABILITY OF DATA AND MATERIALS

The datasets generated and/or analyzed during the current study are available from the corresponding author on reasonable request.

### AUTHOR’S CONTRIBUTIONS

MC conceptualized and designed the study, planned and performed the experiments, preprocessed, analyzed and interpreted the data, and wrote the manuscript. JSG handled the raw Barnes Maze video recordings and developed a pipeline for preprocessing the resulting data. MBV, OM and AB contributed to the wet-lab experimental procedures and imaging. OM and AB performed the quantification of all immunohistochemical stainings. ER was involved in the preprocessing of behavioral data. LLF was involved in maternal scoring and wet-lab work. NR contributed to the analysis and interpretation of microglial morphology. LG provided laboratory facilities, protocols, and guidance for the measurement of soluble and insoluble amyloid. JSG, NR, LG, PJL, RO and HK contributed to the critical revision of the manuscript. PJL, RO, and HK contributed to the conceptualization of the study and provided supervision and guidance throughout the project. All authors read and approved the final manuscript.

## ACKNOWLEDGEMENTS

We sincerely thank Helmut Kessels for his support and for providing resources that contributed to this work. We also thank Magalie Mathias for her valuable technical assistance and support during the experimental work conducted at the IGF, University of Montpellier, France. Finally, we gratefully acknowledge the animals whose use made this research possible.

## Abbreviations

Aβ: amyloid-beta
AD: Alzheimer’s disease
WT: Wild-type
APP/PS1: APPswe/PS1dE9
EH: Early handling
Ctrl: Control
ROI: region of interest
ORT: Novel object recognition test
DI: Discrimination index
NO: Novel object
FO: Familiar object
BM: Barnes maze
HIP: Hippocampal composite
PFC: prefrontal cortex composite
DG: Dentate gyrus
CA1: cornu ammonis 1
PLA: prelimbic area
ACC: anterior cingulate cortex
ILA: infralimbic area

## REFERENCES

[1] 2025 Alzheimer’s disease facts and figures. Alzheimers Dement 2025;21:e70235. 10.1002/alz.70235.

[2] De Strooper B, Karran E. The Cellular Phase of Alzheimer’s Disease. Cell 2016;164:603–15. 10.1016/j.cell.2015.12.056.

[3] Lo Vecchio F, La Torre A, Gravina C, D’Onofrio G, Greco A. Molecular Mechanisms Underlying Alzheimer’s Disease Pathogenesis: Comprehensive Overview. Int J Mol Sci 2026;27:4651. 10.3390/ijms27104651.

[4] Ramachandran AK, Das S, Joseph A, Shenoy GG, Alex AT, Mudgal J. Neurodegenerative Pathways in Alzheimer’s Disease: A Review. Curr Neuropharmacol 2021;19:679–92. 10.2174/1570159X18666200807130637.

[5] Testo AA, Roundy G, Dumas JA. Cognitive Decline in Alzheimer’s Disease. In: Kidd EJ, Newhouse PA, editors. Neurobiol. Alzheimers Dis., vol. 69, Cham: Springer Nature Switzerland; 2024, p. 181–95. 10.1007/7854_2024_527.

[6] Stern Y. Cognitive reserve in ageing and Alzheimer’s disease. Lancet Neurol 2012;11:1006–12. 10.1016/S1474-4422(12)70191-6.

[7] Carrigan M, Bocancea DI, Vogel J, Van Loenhoud AC, Tesi N, Barkhof F, et al. A novel approach to resilience and its links with education and Alzheimer’s disease genetics. Alzheimers Dement 2025;21:e70379. 10.1002/alz.70379.

[8] Lesuis SL, Van Hoek BACE, Lucassen PJ, Krugers HJ. Early postnatal handling reduces hippocampal amyloid plaque formation and enhances cognitive performance in APPswe/PS1dE9 mice at middle age. Neurobiol Learn Mem 2017;144:27–35. 10.1016/j.nlm.2017.05.016.

[9] Webster SJ, Bachstetter AD, Van Eldik LJ. Comprehensive behavioral characterization of an APP/PS-1 double knock-in mouse model of Alzheimer’s disease. Alzheimers Res Ther 2013;5:28. 10.1186/alzrt182.

[10] Lesuis SL, Maurin H, Borghgraef P, Lucassen PJ, Leuven FV, Krugers HJ. Positive and negative early life experiences differentially modulate long term survival and amyloid protein levels in a mouse model of Alzheimer’s disease. Oncotarget 2016;7:39118–35. 10.18632/oncotarget.9776.

[11] García-Domínguez M. Glial Cell Dynamics in Neuroinflammation: Mechanisms, Interactions, and Therapeutic Implications. Biomedicines 2026;14:115. 10.3390/biomedicines14010115.

[12] Verkhratsky A, Butt A, Li B, Illes P, Zorec R, Semyanov A, et al. Astrocytes in human central nervous system diseases: a frontier for new therapies. Signal Transduct Target Ther 2023;8:396. 10.1038/s41392-023-01628-9.

[13] Wang L, Ling H, He H, Hu N, Xiao L, Zhang Y, et al. Dysfunctional synaptic pruning by microglia correlates with cognitive impairment in sleep-deprived mice: Involvement of CX3CR1 signaling. Neurobiol Stress 2023;25:100553. 10.1016/j.ynstr.2023.100553.

[14] Lu A, Chen W-T, Dalby M, Sainz Garcia D, Vanheusden M, de Vries LE, et al. Human microglial transitions at the Aβ-tau inflection point associate with divergent pathways to dementia and resilience. Nat Med 2026;32:2047–59. 10.1038/s41591-026-04393-8.

[15] Baligács N, Albertini G, Borrie SC, Serneels L, Pridans C, Balusu S, et al. Homeostatic microglia initially seed and activated microglia later reshape amyloid plaques in Alzheimer’s Disease. Nat Commun 2024;15:10634. 10.1038/s41467-024-54779-w.

[16] Heneka MT, van der Flier WM, Jessen F, Hoozemanns J, Thal DR, Boche D, et al. Neuroinflammation in Alzheimer disease. Nat Rev Immunol 2025;25:321–52. 10.1038/s41577-024-01104-7.

[17] Oshima T, Kater MSJ, Huffels CFM, Wesseling EM, Middeldorp J, Hol EM, et al. Early amyloid-induced changes in microglia gene expression in male APP/PS1 mice. J Neurosci Res 2024;102:e25295. 10.1002/jnr.25295.

[18] Lee H-G, Lee J-H, Flausino LE, Quintana FJ. Neuroinflammation: An astrocyte perspective. Sci Transl Med 2023;15:eadi7828. 10.1126/scitranslmed.adi7828.

[19] Abu Raya M, Zeltzer E, Blazhenets G, Schonhaut DR, Allen IE, Carrillo MC, et al. Sex differences in amyloid PET in a large, real world sample from the Imaging Dementia– Evidence for Amyloid Scanning (IDEAS) Study. Alzheimers Dement 2025;21:e70304. 10.1002/alz.70304.

[20] Lin KA, Choudhury KR, Rathakrishnan BG, Marks DM, Petrella JR, Doraiswamy PM, et al. Marked gender differences in progression of mild cognitive impairment over 8 years. Alzheimers Dement Transl Res Clin Interv 2015;1:103–10. 10.1016/j.trci.2015.07.001.

[21] Emrani S, Sundermann EE. Sex/gender differences in the clinical trajectory of Alzheimer’s disease: Insights into diagnosis and cognitive reserve. Front Neuroendocrinol 2025;77:101184. 10.1016/j.yfrne.2025.101184.

[22] Sharma S, Gibbons A, Saphire EO. Sex differences in tissue-specific immunity and immunology. Science 2025;389:599–603. 10.1126/science.adx4381.

[23] Klein SL. The effects of hormones on sex differences in infection: from genes to behavior. Neurosci Biobehav Rev 2000;24:627–38. 10.1016/S0149-7634(00)00027-0.

[24] Osborne BF, Turano A, Schwarz JM. Sex differences in the neuroimmune system. Curr Opin Behav Sci 2018;23:118–23. 10.1016/j.cobeha.2018.05.007.

[25] Arenaza Urquijo EM, Boyle R, Casaletto K, Anstey KJ, Vila Castelar C, Colverson A, et al. Sex and gender differences in cognitive resilience to aging and Alzheimer’s disease. Alzheimers Dement 2024;20:5695–719. 10.1002/alz.13844.

[26] Seibenhener ML, Wooten MC. Use of the Open Field Maze to Measure Locomotor and Anxiety-like Behavior in Mice. J Vis Exp 2015:52434. 10.3791/52434.

[27] Lueptow LM. Novel Object Recognition Test for the Investigation of Learning and Memory in Mice. J Vis Exp 2017:55718. 10.3791/55718.

[28] Pitts M. Barnes Maze Procedure for Spatial Learning and Memory in Mice. BIO-Protoc 2018;8. 10.21769/BioProtoc.2744.

[29] Hoeijmakers L, Amelianchik A, Verhaag F, Kotah J, Lucassen PJ, Korosi A. Early-Life Stress Does Not Aggravate Spatial Memory or the Process of Hippocampal Neurogenesis in Adult and Middle-Aged APP/PS1 Mice. Front Aging Neurosci 2018;10:61. 10.3389/fnagi.2018.00061.

[30] Roberts H, Fang Y, Quinn K, Hill T, Peck MR, Bartke A, et al. Lifespan of male and female APP/PS1 and APP^NL-F/NL-F^ mouse models of Alzheimer’s disease 2024. 10.1101/2024.10.15.618508.

[31] Klakotskaia D, Agca C, Richardson RA, Stopa EG, Schachtman TR, Agca Y. Memory deficiency, cerebral amyloid angiopathy, and amyloid-β plaques in APP+PS1 double transgenic rat model of Alzheimer’s disease. PLOS ONE 2018;13:e0195469. 10.1371/journal.pone.0195469.

[32] Torres L, Danver J, Ji K, Miyauchi JT, Chen D, Anderson ME, et al. Dynamic microglial modulation of spatial learning and social behavior. Brain Behav Immun 2016;55:6–16. 10.1016/j.bbi.2015.09.001.

[33] Zong T, Li N, Han F, Liu J, Deng M, Li V, et al. Microglial depletion rescues spatial memory impairment caused by LPS administration in adult mice. PeerJ 2024;12:e18552. 10.7717/peerj.18552.

[34] León-Rodríguez A, Fernández-Arjona MDM, Grondona JM, Pedraza C, López-Ávalos MD. Anxiety-like behavior and microglial activation in the amygdala after acute neuroinflammation induced by microbial neuraminidase. Sci Rep 2022;12:11581. 10.1038/s41598-022-15617-5.

[35] Zheng Z-H, Tu J-L, Li X-H, Hua Q, Liu W-Z, Liu Y, et al. Neuroinflammation induces anxiety- and depressive-like behavior by modulating neuronal plasticity in the basolateral amygdala. Brain Behav Immun 2021;91:505–18. 10.1016/j.bbi.2020.11.007.

[36] Davies DS, Ma J, Jegathees T, Goldsbury C. Microglia show altered morphology and reduced arborization in human brain during aging and A lzheimer’s disease. Brain Pathol 2017;27:795–808. 10.1111/bpa.12456.

[37] Paasila PJ, Davies DS, Kril JJ, Goldsbury C, Sutherland GT. The relationship between the morphological subtypes of microglia and Alzheimer’s disease neuropathology. Brain Pathol 2019;29:726–40. 10.1111/bpa.12717.

[38] Ali S, Liu X, Queen NJ, Patel RS, Wilkins RK, Mo X, et al. Long-term environmental enrichment affects microglial morphology in middle age mice. Aging 2019;11:2388–402. 10.18632/aging.101923.

[39] Stuart KE, King AE, King NE, Collins JM, Vickers JC, Ziebell JM. Late-life environmental enrichment preserves short-term memory and may attenuate microglia in male APP/PS1 mice. Neuroscience 2019;408:282–92. 10.1016/j.neuroscience.2019.04.015.

[40] Chen H, Cao J, Zhang Y, Su X, Kang Y, Gao L, et al. Enriched Environment Alleviate AD Pathological Progression by Reducing Microglia Complement Signaling in Aged Male APP / PS1 Mice. FASEB J 2025;39:e71026. 10.1096/fj.202501545RR.

[41] Chin J, Scharfman HE. Shared cognitive and behavioral impairments in epilepsy and Alzheimer’s disease and potential underlying mechanisms. Epilepsy Behav 2013;26:343–51. 10.1016/j.yebeh.2012.11.040.

[42] Vossel KA, Ranasinghe KG, Beagle AJ, Mizuiri D, Honma SM, Dowling AF, et al. Incidence and impact of subclinical epileptiform activity in Alzheimer’s disease. Ann Neurol 2016;80:858–70. 10.1002/ana.24794.

[43] Minkeviciene R, Rheims S, Dobszay MB, Zilberter M, Hartikainen J, Fülöp L, et al. Amyloid β-Induced Neuronal Hyperexcitability Triggers Progressive Epilepsy. J Neurosci 2009;29:3453–62. 10.1523/JNEUROSCI.5215-08.2009.

[44] Benjamin DJ, Berger JO, Johannesson M, Nosek BA, Wagenmakers E-J, Berk R, et al. Redefine statistical significance. Nat Hum Behav 2017;2:6–10. 10.1038/s41562-017-0189-z.

