## Supplementary for "The modulatory role of early-life experiences and sex on cognitive resilience in APP/PS1 mice over time"

**Supplementary tables (ST)**

**ST1**. Sholl analysis-derived AUC (intersections·µm) in APP/PS1: PAM vs NPAM

| Age | Brain region | F-value | df | p-value | PAM | NPAM |
| --- | --- | --- | --- | --- | --- | --- |
| 6 months | HIP | 28.05 | 1, 506.78 | <0.001 | 72 ± 75 | 212 ± 90 |
|  | PFC | 13.88 | 1, 331.31 | <0.001 | 98 ± 100 | 219 ± 91 |
|  | DG | 17.08 | 1, 250.64 | <0.001 | 84 ± 80 | 198 ± 89 |
|  | CA1 | 14.35 | 1, 250.41 | <0.001 | 31 ± 29 | 225 ± 89 |
|  | ACC | 5.33 | 1, 103.74 | <0.05 | 120 ± 119 | 205 ± 87 |
|  | PLA | 6.55 | 1, 98.66 | <0.05 | 80 ± 92 | 228 ± 98 |
|  | ILA | 6.90 | 1, 108 | <0.01 | 57 ± 32 | 224 ± 86 |
| 12 months | HIP | 117.72 | 1, 599.39 | <0.001 | 78 ± 49 | 148 ± 76 |
|  | PFC | 71.63 | 1, 424.44 | <0.001 | 72 ± 53 | 134 ± 72 |
|  | DG | 42.68 | 1, 262.44 | <0.001 | 77 ± 50 | 127 ± 64 |
|  | CA1 | 38.11 | 1, 289.29 | <0.001 | 81 ± 47 | 162 ± 81 |
|  | ACC | 24.29 | 1, 134.71 | <0.001 | 68 ± 38 | 121 ± 38 |
|  | PLA | 24.29 | 1, 135.29 | <0.001 | 72 ± 63 | 136 ± 66 |
|  | ILA | 15.02 | 1, 134.87 | <0.001 | 81 ± 62 | 144 ± 78 |

**ST2**. Microglia HCPC distribution in APP/PS1: PAM vs NPAM

| Age | Brain region | F-value | df | p-value |
| --- | --- | --- | --- | --- |
| 6 months | HIP | 27.12 | 1, 32 | <0.001 |
|  | PFC | 13.56 | 1, 23 | <0.001 |
|  | DG | 14.28 | 1, 31 | <0.001 |
|  | CA1 | 18.54 | 1, 25 | <0.001 |
|  | ACC | 6.48 | 1, 21 | <0.001 |
|  | PLA | 4.17 | 1, 20 | <0.01 |
|  | ILA | 3.45 | 1, 19 | <0.05 |
| 12 months | HIP | 21.87 | 1, 53 | <0.001 |
|  | PFC | 18.33 | 1, 49 | <0.001 |
|  | DG | 10.38 | 1, 51 | <0.001 |
|  | CA1 | 10.71 | 1, 45 | <0.001 |
|  | ACC | 5.27 | 1, 44 | <0.001 |
|  | PLA | 5.88 | 1, 43 | <0.001 |
|  | ILA | 9.52 | 1, 36 | <0.001 |

**Supplementary figures (SF)**

**SF1**. Survival of EH vs Ctrl offspring (main cohort)

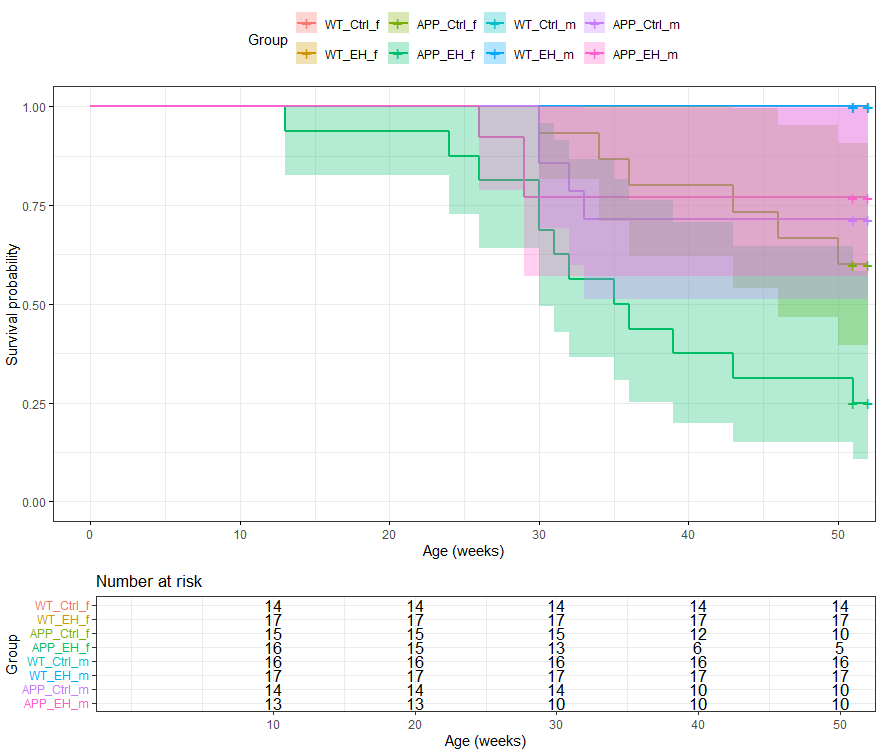

**SF2. Markers of pathology in APP/PS1 mice by condition**

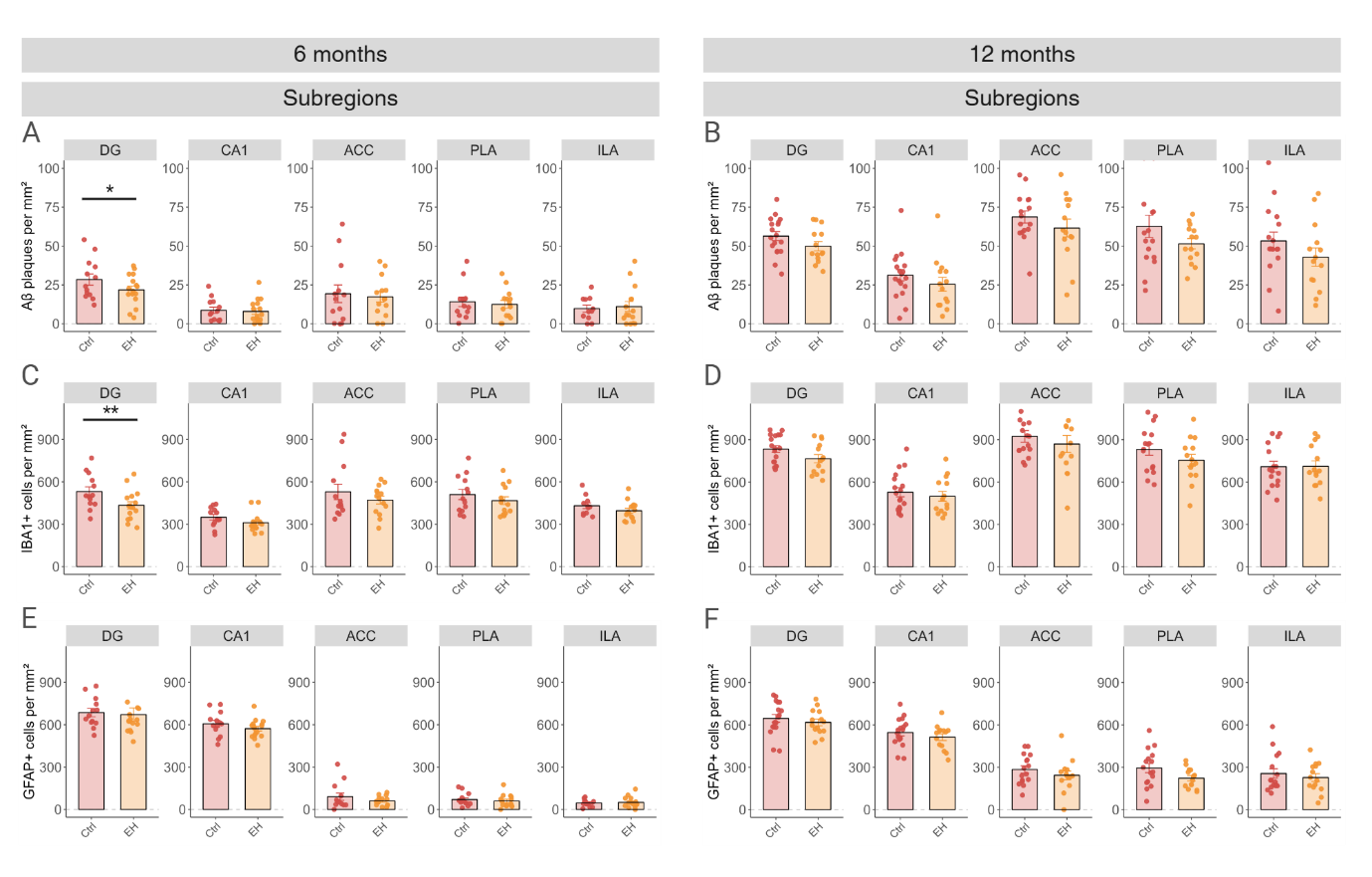

**SF3. Markers of pathology in APP/PS1 mice by sex**

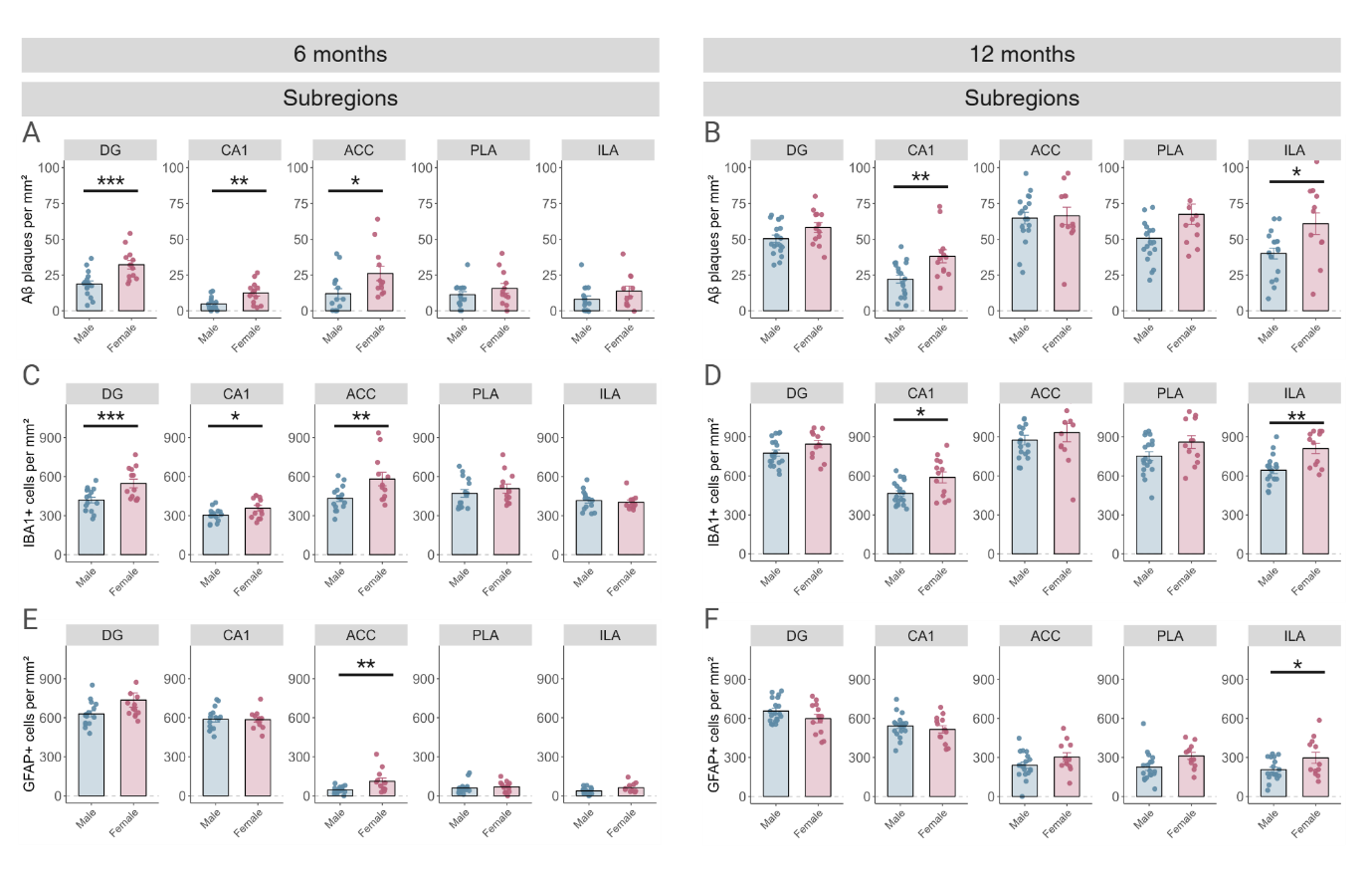

**SF4. Behavioral outcomes in WT mice**

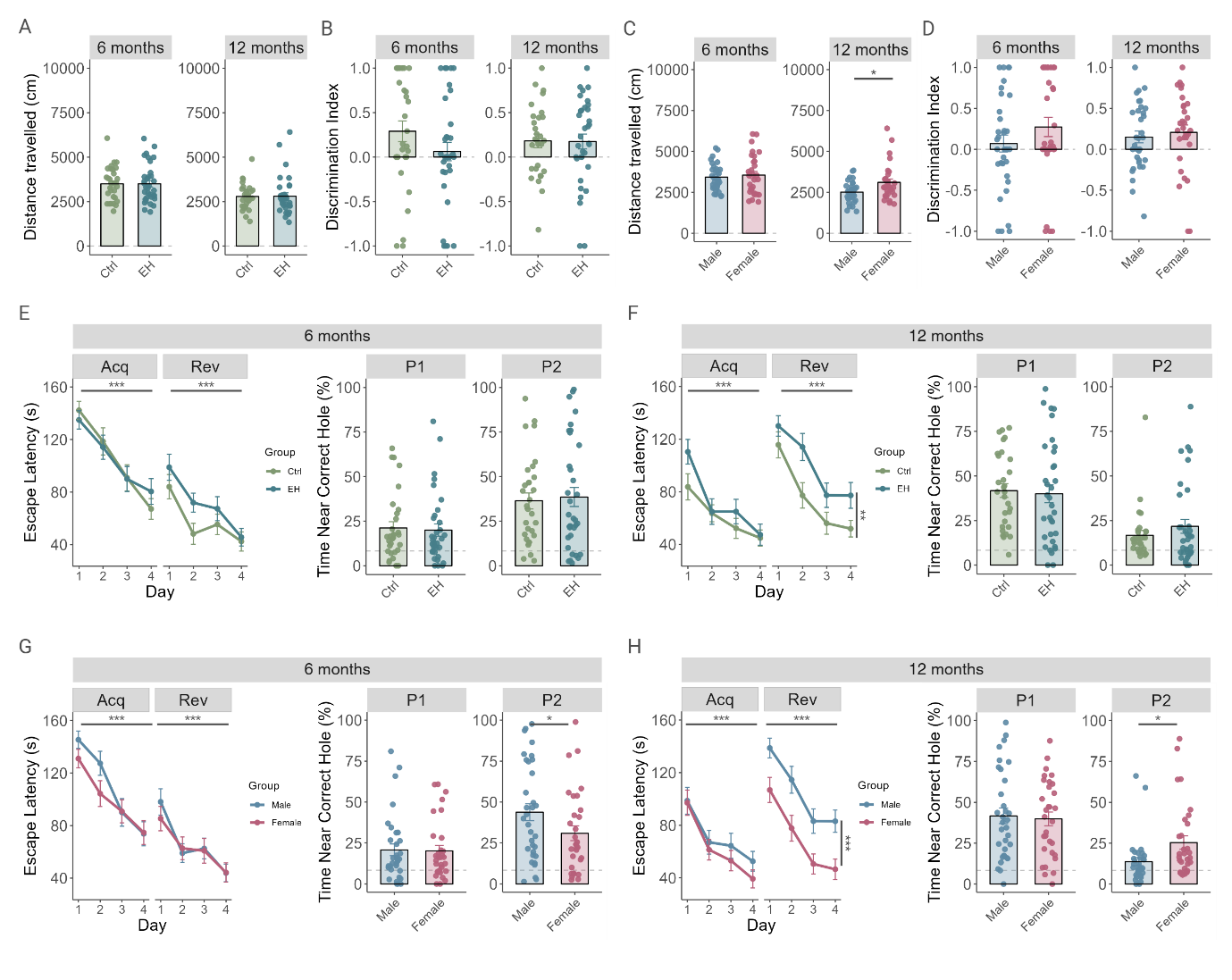

**SF5. Markers of pathology in WT mice**

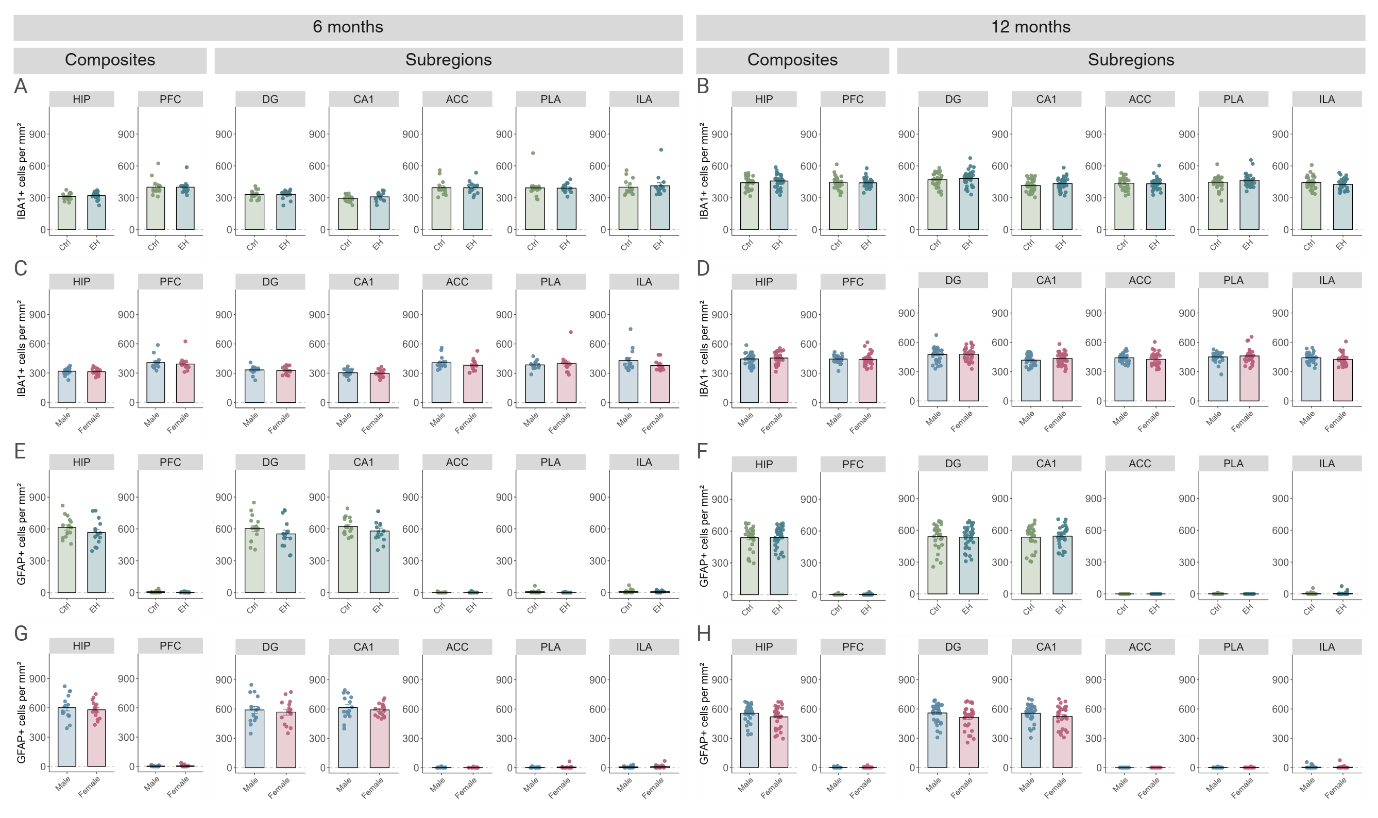

**SF6. Associations of cognition and pathology in 12-month-old APP/PS1 mice**

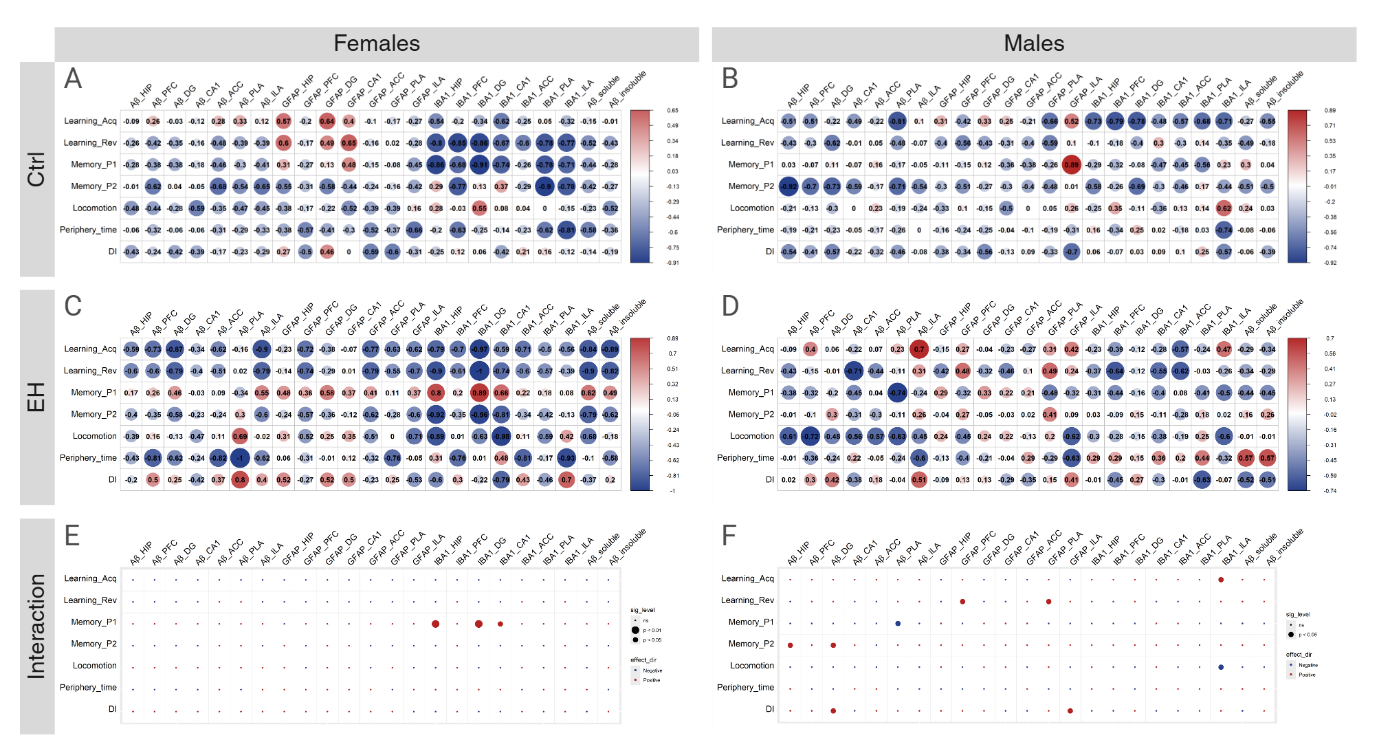

**SF7. Microglia morphology in subregions of 6- and 12-month-old APP/PS1 mice**

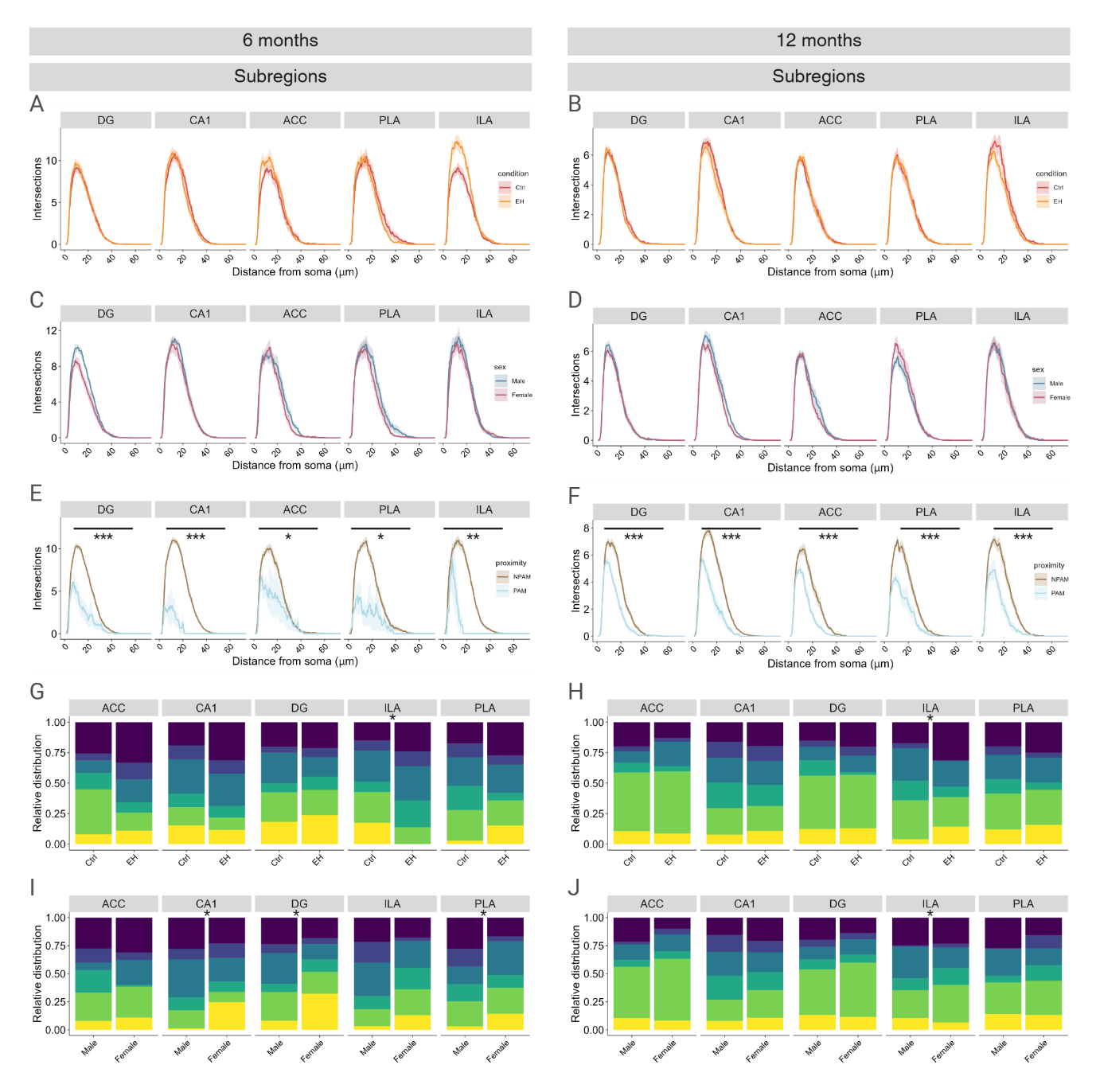

**SF8. Microglia morphology in 6- and 12-month-old WT mice**

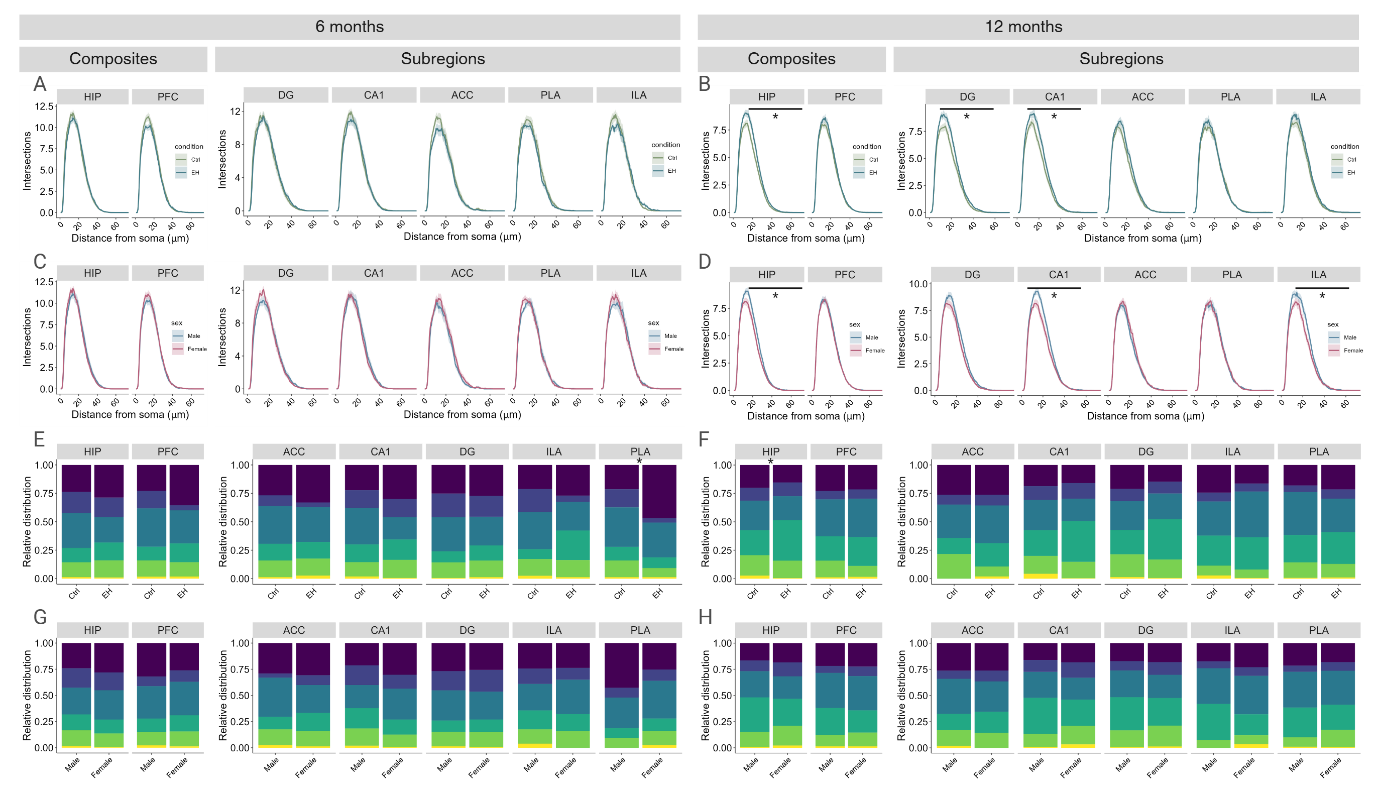

**Supplementary analyses (SA)**

**SA1. Survival analysis of EH vs Ctrl offspring (main cohort)**

Survival was visualized using Kaplan–Meier curves compared between groups using the log-rank (Mantel–Cox) test. Survival differed significantly across the eight Genotype × Condition × Sex groups (χ²=60.5, p<0.001). Survival analysis using firth penalized Cox regression on genotype revealed that APP/PS1 mice had a significantly higher hazard of death compared to WT (HR=70.98, 95%CI [10.06, 9134.83], p<0.001). Due to no events in WT, we tested effects of condition and sex in APP/PS1 animals only and found that EH had a significantly higher hazard of death compared to Ctrl (HR=2.66, 95%CI [1.06, 7.38], p<0.05), whereas sex was not significantly associated with survival (p=0.70), and there were no significant interactions (p=0.19).

**SA2. Maternal Care**

Group differences between EH and control were assessed using Welch’s t-tests across genotypes. For variables with clear deviations from normality (harmful contact [HC], passive/blanket nursing [PBN]), results were confirmed using Wilcoxon rank-sum tests. Contact was combined into a net Contact score: net Contact = Contact – (nest exit + no Contact). EH dams showed increased maternal behavior across active measures including arched-back nursing (ABN, t=-2.29, p<0.05), licking/grooming (LG, t=-2.24, p<0.05) and nest building (NB, t=-4.11, p<0.001), whereas control dams showed higher net contact with their pups (t=7.72, p<0.001) during the scoring. There was no difference between dams in PBN (t=0.15, p=0.88) or harmful contact (HC, t=-1.81, p=0.08). Nest visibility was higher in EH dams compared to control dams (t=-4.18, p<0.001).

**SA3. Body weight pups**

Group differences between EH and control were assessed using Welch’s t-tests across genotypes. Because individual pup identities could not be tracked across P2-P9, body weight gain from P2 to P9 was calculated at the litter level per sex by subtracting the mean P2 weight from the mean P9 weight for each litter. There was no difference in body weight gain between EH and control litters (t=1.43, p=0.16) in the pooled-sex, nor in a sex-stratified analysis (f: t=1.12, p=0.27 and m: t=0.91, p=0.37).

**SA4. Open Field Test**

Overall, APP/PS1 mice displayed increased locomotor activity compared to WT mice at 6 months of age (genotype effect: F(1,112) = 36.14, p < 0.001), consistent with less periphery time compared to WT mice (genotype effect: F(1,112) = 5.05, p < 0.05). At 12 months, APP/PS1 mice remained more active than WT animals (genotype effect: F(1,87) = 6.64, p < 0.05), albeit with no difference in anxiety-like behavior (genotype effect: F(1,87) = 1.35, p = 0.25). Among WT animals only, we found that while WT showed a similar sex effect at this age (F(1,58) = 6.97, p < 0.05), there was no comparable condition effect (F(1,58) = 0.01, p = 0.93), nor interaction (p = 0.71), indicating that EH effects on general locomotion at 12 months are APP/PS1-specific.

**SA5. Barnes maze FET at 12 months**

To test whether mice had a long-term recall at 12 months of the last escape hole location, we treated the FET at 12 months as an additional probe trial prior to acquisition training. While WT-Ctrl animals correctly identified the last escape hole location trained at 6 months of age (t(29) = 2.42, p < 0.05), we found no recall of this location in APP/PS1 mice (APP-Ctrl: t(18) = 0.02, p = 0.10; APP-EH: t(13) = 0.71, p = 0.49), nor in WT-EH animals (t(33) = 1.20, p = 0.24).
